# Diverse conjugative and mobilisable elements underpin key adaptive traits in *Xanthomonas*

**DOI:** 10.64898/2026.06.24.734381

**Authors:** Elena Colombi, Timothy M. Ghaly, Nishadi Samarakoon, Vaheesan Rajabal, Sasha G. Tetu

## Abstract

Horizontal gene transfer mediated by mobile genetic elements (MGEs) is a major driver of bacterial evolution and ecological adaptation. In the plant-associated genus *Xanthomonas*, multiple MGEs have been implicated in virulence, host specialisation, and environmental persistence, yet MGE diversity and evolutionary dynamics across the genus remain poorly understood. Here, we performed a comparative analysis of conjugative and mobilisable plasmids, integrative and conjugative elements (ICEs), integrative and mobilisable elements (IMEs), and their cargo genes across 516 complete genomes of three major *Xanthomonas* species: *X. campestris*, *X. cissicola*, and *X. oryzae*. We identified pronounced interspecific differences, with *X. cissicola* and *X. campestris* harbouring large and diverse MGE repertoires, comprising 28.3% and 26.7% of their respective pangenomes, whereas *X. oryzae* contained far fewer MGEs, making up only 3.6% of the identified pangenome. These differences were associated with host defence systems, including CRISPR-Cas and restriction-modification systems, and with variation in CRISPR spacer diversity. IMEs were the most abundant MGEs across all species, encoding diverse defence systems and accessory genes. ICEs exhibited signatures of horizontal transfer within and between species, and across genera. Notably, nearly identical ICEs carrying heavy-metal resistance genes were identified in *Xanthomonas* and *Pseudomonas aeruginosa*, indicating recent transfer between genera. MGEs collectively carried genes involved in virulence, interbacterial interactions, defence against phages, and plant cell wall degradation, with several elements associated with specific pathovars. Together, our findings establish MGEs as key drivers of genome plasticity and adaptive evolution in *Xanthomonas*, shaped by a dynamic interplay with host defence systems.

## INTRODUCTION

Horizontal gene transfer (HGT) is a major driver of bacterial evolution, influencing ecological adaptation, and diversification [1–4]. By enabling the transfer of suites of genes that have already been refined by selection in a donor lineage, HGT can rapidly confer complex traits to recipient bacteria, allowing colonization of new ecological niches in a single evolutionary step [4–7]. Mobile genetic elements (MGEs) are key mediators of HGT [8]. Conjugative plasmids and integrative and conjugative elements (ICEs) are self-transmissible MGEs, encoding the conjugation machinery required for their own transfer between bacterial cells. In contrast, mobilizable plasmids and integrative and mobilizable elements (IMEs) lack a complete conjugation system but can exploit the conjugative apparatus of co-resident plasmids or ICEs through their encoded relaxase proteins. In plant-associated bacteria, such MGEs play a central role in adaptive evolution by promoting the horizontal acquisition of virulence determinants [9–11], antimicrobial and heavy-metal resistance genes [12–15], and metabolic functions [16–18]. Through these processes, MGEs contribute to host specialization, expansion of host range, and persistence in diverse agricultural and environmental settings, shaping the evolutionary trajectories of phytopathogens.

*Xanthomonas* is a genus of Gram-negative bacteria in the class Gammaproteobacteria that contains pathogenic strains able to cause disease in more than 400 different plants. Bacteria of the genus *Xanthomonas* affect several crops of economic importance, responsible for diseases such as bacterial black spot (*X. campestris*), citrus canker (*X. citri/X. cissicola*), and the bacterial leaf blight in rice (*X. oryzae*) [19]. In common with other genera of plant pathogens, *Xanthomonas* also contains strains associated with plants that do not cause disease symptoms [20–22].

In *Xanthomonas*, both ICEs and plasmids have been shown to play pivotal roles in genome plasticity, environmental adaptation, and the emergence of epidemic lineages. In *X. arboricola* pv. *juglandis*, an ICE carrying copper-resistance genes confers phenotypic resistance to copper-based compounds commonly used in agriculture [23], and notably, this ICE has been detected in strains associated with walnut blight epidemics in France [24]. Plasmids represent another major source of genomic variability in *Xanthomonas* and have been repeatedly implicated in the diversification of virulence repertoires. For instance, a large plasmid was described in *X. citri* (*cissicola*) pv. *citri*, harbouring multiple copies of transcription activator-like effector (TALE) genes within a single replicon [25], and in *X. citri* (*cissicola*) pv. *viticola*, curing of native plasmids resulted in a marked reduction in exopolysaccharide production and virulence [26].

Despite their recognized importance, MGEs in *Xanthomonas* have so far been characterized mainly in individual strains, epidemic lineages, or specific pathovars, often in the context of particular traits such as copper resistance or virulence. A comprehensive, comparative view of MGE diversity, distribution, and evolutionary dynamics across the genus has been lacking. Here, we address this gap by systematically analysing conjugative and mobilizable plasmids, ICEs, and IMEs across the three *Xanthomonas* species (*X. oryzae*, *X. campestris*, and *X. cissicola*), which represent the three Xanthomonas species with the highest number of complete genomes in public databases. By leveraging large, taxonomically diverse genome collections, our aims to explore the landscape of MGEs in *Xanthomonas* and to assess their contribution to genome evolution, virulence, and host specificity.

## METHODS

### *Xanthomonas* classification and phylogeny

Complete genomes of *Xanthomonas* were downloaded from the National Center for Biotechnology Information (NCBI) Database (NCBI txid: 338) (last accessed January 2025). Taxonomic classifications of the genomes was based on the Genome Taxonomy Database (GTDB) [27] release 2.2.0 using GTDB-Tk v2.4.0 [28]. The command classify_wf was used with default settings. GTDB-Tk first aligns 120 single-copy phylogenetic marker genes, then classifies each genome based on its placement into domain-specific reference trees (built from 47,899 prokaryote genomes), its relative evolutionary divergence, and average nucleotide identity to reference genomes in the GTDB. Genomes of the species that had more than 100 genomes in the database (*X. campestris* (n=205), *X. oryzae* (n=187), and *X. cissicola* (n=124)) (Table S1) were selected and annotated with Bakta [29].

Species phylogeny was built using Realphy [30] which maps genomes to a series of reference genomes via bowtie2 2.0.0-beta7 [31]. From these, multiple sequence alignments are constructed and used to infer phylogenetic trees via PhyML 3.1 [31]. Trees are visualised using the ggtree 3.16 [32] and ggtreeExtra 2.3 [33] R packages.

### Mobile genetic element search and annotation

Conjugative DNA relaxases, also referred to as ‘Mob’ proteins, are DNA endonucleases that target and nick the origin-of-transfer sequences present on MGEs. Relaxases are required for recruitment of ssDNA of conjugative and mobilizable elements to the conjugative type IV secretion system (T4SS) for transfer. Relaxase proteins have been assigned into six MOB families: MOBF, MOBH, MOBQ, MOBC, MOBP and MOBV [34]. To identify putative horizontally transmissible elements, relaxase gene sequences were searched with hmmscan from HMMER [35] using available hidden Markov model (HMM) protein profiles (MOB database) [36] defined for distinct MOB-protein families [34]. For this search, a bit score threshold (-T) of 33 was used. For ambiguous classifications, the MOB-protein family was chosen by the best e-value in the full sequence. To delineate the chromosomal boundaries of identified ICEs and IMEs, sequence regions surrounding the MOB genes and/or conjugation-gene clusters were inspected for integrase genes and flanking DNA sequence repeats commonly present within integrase attachment (*att*) sites. Elements lacking identifiable *att* sites were excluded (Table S2).

A subset of non-redundant MGEs (Table S3) was identified from the initial MGEs based on their average nucleotide identity (ANI) (ANI > 98% and coverage > 98%), and on differences in gene content within ANI clusters. ANI was calculated with FastANI 1.33 [37] run with ––fragLen 1000 –s. MGEs were considered of the same non-redundant group if they differed by transposases or insertion sequences (ISs) insertions. Fasta and Bakta annotated sequences were deposited at https://github.com/EC-MQ/Xanthomonas_MGEs.

We used multiple approaches to annotate the MGEs-encoded proteins. Proteins were functionally annotated with eggNOG-mapper v2 [38, 39], executed in DIAMOND [40].

To detect genes encoding type III secretion effectors (T3SE), we used a database of 66 previously characterized 66 *Xanthomonas* T3SE retrieved from the EuroXanth platform [41] [42]. T3SEs were identified with Proteinortho v6.0.22 [43] by querying the curated effectors database against the amino acid sequences of the cassette-encoded genes.

DefenseFinder [44] was used to detect known anti-phage systems. The number of spacer in CRISPR-Cas arrays was calculated CRISPRCasFinder [45]. Statistical analyses were executed in R (v4.3). To compare the distribution of MGE abundance between CRISPR-positive and CRISPR-negative genomes, non-parametric analysis was conducted using the two-tailed Mann-Whitney U test. To model MGE counts as a function of CRISPR status, generalized linear models were constructed. Initial models were fitted using a standard Poisson distribution.

Pan-genome association analysis was conducted with Roary [46] and Scoary2 [47] using default setting, Scoary2 was run with ––n-permut 1000. Significant associated genes were selected if the empirical_p values was <0.02 and if the number of strains that have the gene was more than the number of strains that didn’t have the gene had the trait (g+t+>g-t+) and more than the number of strains that had the gene but not the trait (g+t+>g+t-).

### ICE phylogeny and phylogenetic network analysis

To create ICE phylogeny, Proteinortho v6.0.22 [43] was run with default settings to identify single-copy core genes. The nucleotide sequences of single copy orthologues were then aligned with the PRANK algorithm [48], and core gene alignments were concatenated and stripped of gaps with Goalign (https://github.com/evolbioinfo/goalign). FastTree 2.1 [49] was used to build an approximately-maximum-likelihood phylogenetic tree. The tree was rooted at midpoint using the midpoint() function from the phangorn package in R [51] and then it was visualized using the ggtree 3.16 [32] and ggtreeExtra 2.3 [33] R packages. Similarly, for the phylogenetic network analysis to create the Neighbour-Net tree, Proteinortho v6.0.22 [43] was used to identify single-copy core genes, which were aligned with the PRANK algorithm [48], then concatenated and stripped of gaps with Goalign (https://github.com/evolbioinfo/goalign). The alignment was used for the phylogenetic network analysis conducted with the SplitsTree App [50].

### Network analysis

Orthologous gene clusters among MGEs were identified using ProteinOrtho v6.0.22 [43]. Pairwise similarity in gene content between MGEs was quantified using the Jaccard index, calculated as the ratio of the number of shared orthologous gene clusters to the total number of gene clusters present in either MGE. Jaccard similarity was computed for all pairwise combinations of MGEs based on their binary presence–absence profiles. The resulting similarity values range from 0 (no shared orthologous gene) to 1 (shared orthologous gene). To reduce noise and improve interpretability, edges below Jaccard similarity <0.3 threshold were removed prior to visualization. The filtered MGE similarity network was visualized using Cytoscape [51]. In the gene-content similarity network, the nodes represent MGEs and the edges represent the pairwise Jaccard index. Edge weights were mapped to edge thickness to reflect the degree of shared gene content between MGEs. To identify groups of MGEs with similar gene repertoires, clustering was performed using the Markov Cluster Algorithm (MCL) implemented in the Cytoscape plugin clusterMaker2 [52], with Jaccard similarity values used as edge weights.

## RESULTS

### Landscape of conjugative and mobilizable plasmids, ICEs, and IMEs in major *Xanthomonas* **species.**

To look for conjugative and mobilizable elements, complete genomes of *Xanthomonas campestris* (n=205), *X. oryzae* (n=187), and *X. cissicola* (n=124) were searched for conjugative DNA relaxases (also referred to as ‘Mob’ proteins, which target and nick the origin-of-transfer sequences on MGEs and present them to conjugation systems). Overall, both *X. campestris* and *X. cissicola* contained more genomes harbouring at least one MGE than genomes lacking MGEs altogether, while *X. oryzae* was almost devoid of MGEs (Fig. 1 and 2). On average, *X. cissicola* harboured a greater number of MGEs per genome than *X. campestris*, with maxima of nine and seven MGEs per genome, respectively. IMEs were the most abundant class of MGEs, being present in 80% of *X. campestris* genomes and 87% of *X. cissicola* genomes. In contrast, intact ICEs were detected in 25% and 58% of genomes, conjugative plasmids in 16% and 59%, and mobilizable plasmids in 0% and 44% of *X. campestris* and *X. cissicola* genomes, respectively (Fig. 1 and 2).

**Fig. 1.**
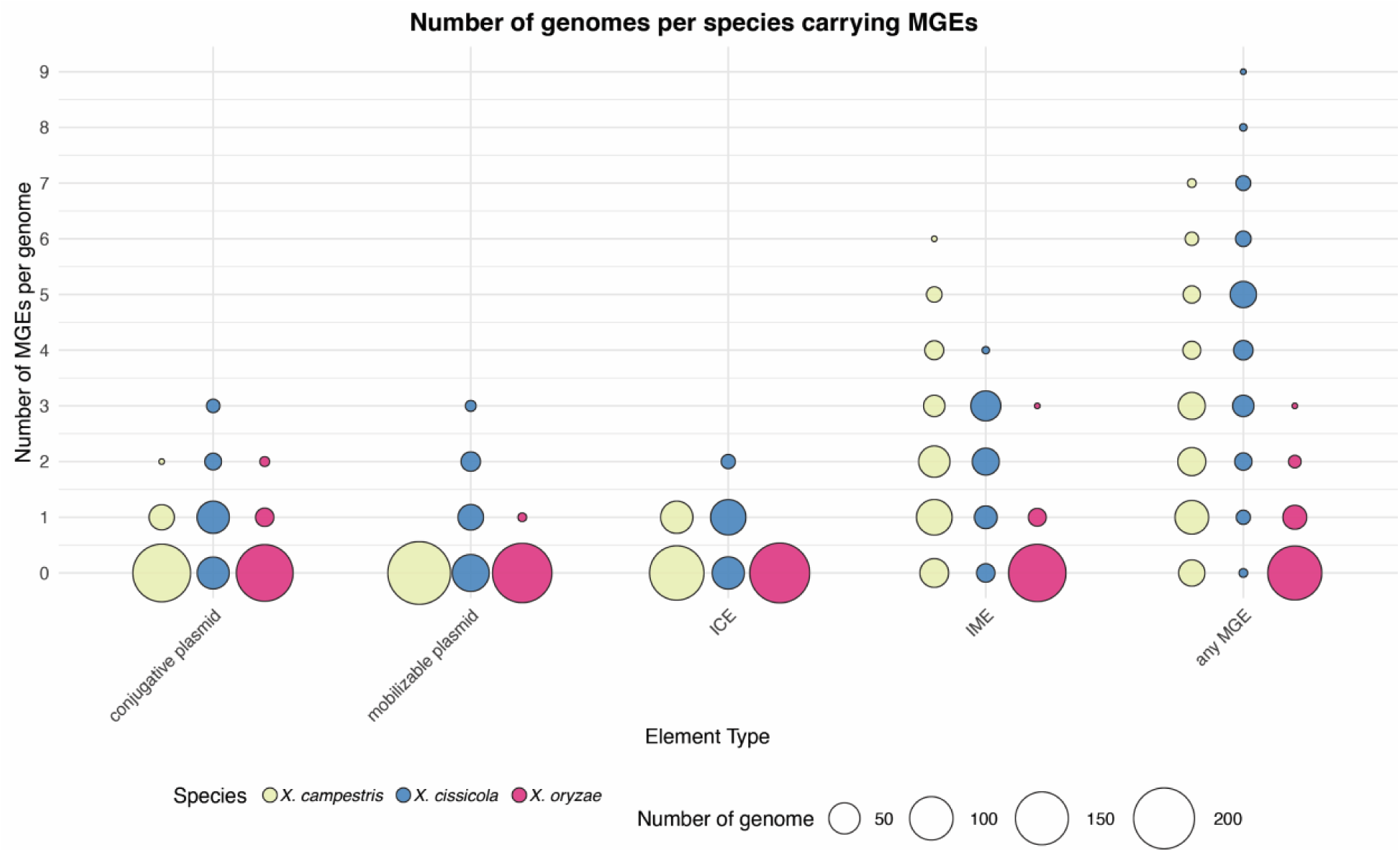
– Distribution of mobile genetic elements across genomes. Bubble plot showing the number of genomes per species carrying different classes of mobile genetic elements (conjugative plasmids, mobilizable plasmids, ICEs, and IMEs). The y-axis indicates the number of elements per genome, circle size reflects the number of genomes, and colors denote species.

**Figure 2.**
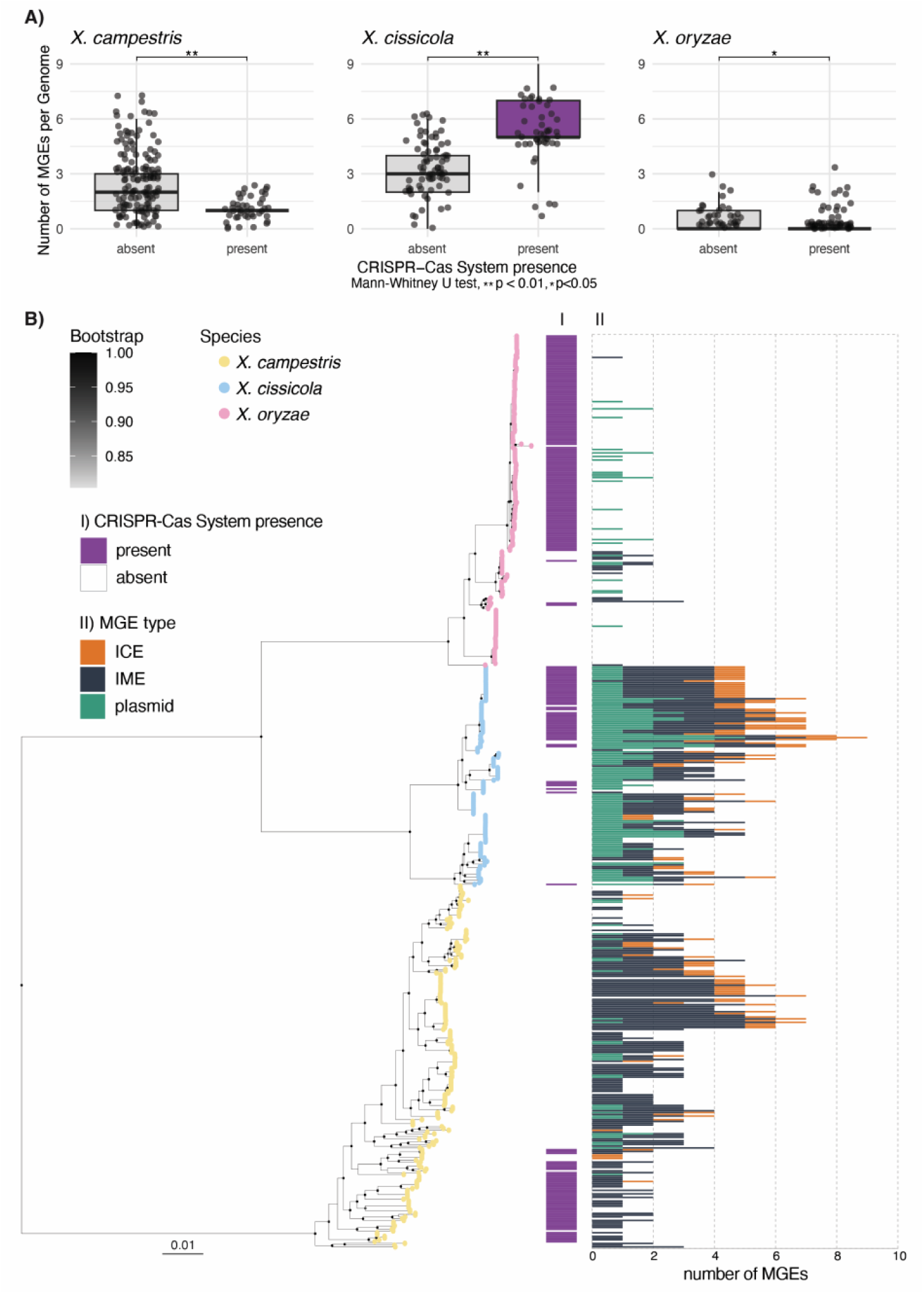
Phylogenetic distribution and association of CRISPR-Cas systems and MGEs. A) Boxplots illustrate the number of MGEs per genome in *X. campestris*, *X. cissicola*, and *X. oryzae*, partitioned by the presence or absence of CRISPR-Cas systems. Individual data points represent individual genomes. Statistical significance was determined using the Mann-Whitney U test (** *p* < 0.01, * *p* < 0.05). B) Core genome phylogeny of *X. campestris*, *X. cissicola*, and *X. oryzae* was built on 515 core genes alignment (256kb). The tree was rooted with *X. translucens* (GCA_000972745) then the tip was removed from the tree. Circles at nodes represent bootstrap support values; only values >80 are shown. Tip labels show species, which were assigned with GTDB-Tk v2 (Table S1). The scale bar indicates substitutions per site. Panel I. Indicates the presence of a CRISPR-Cas system. Panel II. shows the number of MGEs carried by the corresponding isolate.

In total, we identified 33 conjugative plasmids, 0 mobilizable plasmids, 50 ICEs and 356 IMEs in *X. campestris* (with 171 genomes having at least one MGE) (Fig. S1); 107 conjugative plasmids, 77 mobilizable plasmids, 82 ICEs and 239 IMEs in *X. cissicola* (121 genomes with at least one MGE) (Fig. S2); and 24 conjugative plasmids, 3 mobilizable plasmids, 0 ICEs and 18 IMEs in *X. oryzae* (36 genomes with at least one MGE) (Fig. S3) (Table S1 and S2).

The set of non-redundant MGEs was defined from the initial collection based on average nucleotide identity (ANI) and differences in gene content within ANI clusters. Using this approach, we identified as non-redundant MGEs: 27 conjugative plasmids, 16 ICEs, and 115 IMEs in *X. campestris*; 38 conjugative plasmids, 27 mobilizable plasmids, 16 ICEs, and 57 IMEs in *X. cissicola*; and 15 conjugative plasmids, 3 mobilizable plasmids, and 11 IMEs in *X. oryzae*. This set of non-redundant MGEs was used for all subsequent analyses (Table S3).

The distribution of relaxases and T4SS subtypes showed distinct patterns of association based on the type of MGE. In *X. campestris* and *X. cissicola*, ICEs exhibited high conservation, all harboring a MobH relaxase and a Type G (tfc) T4SS. Integration of these ICEs was highly site-specific, primarily occurring adjacent to tRNA-Gly, except for a single ICE integrated adjacent to tRNA-Leu in *X. campestris* and one to *guaA* in *X. cissicola*. The majority of IMEs encoded a MobQ relaxase, but integration occurred across a diverse array of tRNA genes and chromosomal sites (tRNA –Asp, –Arg, –Gly, –Leu, –Ser and –Val, *mutS*, *ssrA*, and an antranilate synthase encoding gene). Conjugative plasmids demonstrated the greatest diversity. While MobP1 and MobF relaxases were predominantly associated with Type T T4SS, we identified instances in *X. oryzae* and *X. cissicola* where these relaxases were coupled with Type I or Type F conjugation systems. Mobilizable plasmids mostly carried a MobF or MobP1 relaxase, or occasionally a MobQ or MobT (Table S2 and S3).

### Presence of CRISPR–Cas systems partially explain differences in MGE content across ***Xanthomonas* species**

Among the species considered, *X. oryzae* harboured a relatively small number of MGEs (total of 45 MGEs in 187 genomes) in comparison to *X. cissicola* (505 MGEs in 124 genomes) and *X. campestris* (439 MGEs in 205 genomes) (Fig. 2). To investigate possible reasons for the marked difference in abundance of MGEs between species, we looked at the prevalence of CRISPR-Cas systems. In *X. oryzae*, the association between CRISPR-Cas presence and MGE count was small but significant (Mann-Whitney U test, p = 0.012). Genomes harbouring CRISPR-Cas systems tended to contain fewer MGEs (mean ≈ 0.21 MGEs) compared to those without CRISPR-Cas (mean ≈ 0.38 MGEs) (Fig. 2A). These results suggest a modest role for CRISPR-Cas in limiting MGE proliferation within *X. oryzae*, with other factors likely also contributing to MGE diversity. Supporting the role of additional defense systems, it was observed that for the African-like strains of *pv. oryzae* which do not harbour a CRISPR-Cas system only one of 31 has an IME while the 122 Asian-like strains of pv. *oryzae*, which generally harbour a CRISPR-Cas system, had 13 non-redundant plasmids and two IMEs (Fig. S3). To identify alternative defence systems in the African-like pv. *oryzae* strains that might explain the very low MGE carriage in this pathovar we performed a pan-genome association analysis [53]. Two defence systems (a type I and a type II restriction-modification systems) were found associated with this pathovar (Bonferroni-corrected p value < 1×10^−14^).

In *X. campestris*, genomes with CRISPR-Cas systems had significantly fewer MGEs (average ≈ 1.00) than those lacking CRISPR-Cas (average ≈ 2.48; Mann-Whitney U test, p < 0.001) (Fig. 2A). To investigate this relationship, a generalized linear model was employed, confirming that CRISPR-Cas presence is a highly significant negative predictor of MGE count (t = –5.56, p < 0.001). The model estimates that the presence of a CRISPR-Cas system in *X. campestris* is associated with an approximately 60% reduction in MGE abundance (IRR ≈ 0.40). In *X. campestris*, the loss of the CRISPR-Cas defence system has been correlated with the diversification of the pv. *Campestris* [54], in fact pv. *campestris* strains harbour 150 of the 179 non-redundant MGEs present in the species.

Curiously, in *X. cissicola*, there was a positive association between CRISPR-Cas presence and MGE abundance (Mann-Whitney U test, p < 0.001). Genomes with CRISPR-Cas systems had significantly higher MGE counts (mean ≈ 5.27) compared to those without (mean ≈ 3.32) (Fig. 2A). To explore this further we looked at the number of spacers found in each CRISPR-Cas array.

*X. cissicola* has on average only 16.6 +/-2.9 whereas *X. oryzae* has 71.6 +/− 19.8 spacers per array while *X. campestris* harbours 92 +/− 28.1 spacers per array. This suggests that factors such as the number of spacers, which reflect the evolutionary history and diversity of the CRISPR-Cas immune repertoire, could influence its effectiveness in controlling MGEs. A higher and more diverse spacer pool, as observed in *X. campestris,* may provide broader protection, leading to a negative correlation between CRISPR-Cas activity and MGE abundance. Conversely, a limited spacer repertoire, such as in *X. cissicola*, might be insufficient for effective defence, potentially explaining instances where there is poor correlation between presence/absence of CRISPR-Cas system and MGE load.

### Inter– and intra-species movement of MGEs in *Xanthomonas*

A gene-content similarity network of the non-redundant MGEs was constructed using the Jaccard index, revealing a structured organization of MGEs into distinct clusters (Fig. 3). Clustering was driven primarily by MGE type rather than host species, with conjugative plasmids, mobilizable plasmids, ICEs and IMEs generally forming separate groups, probably reflecting a different makeup of backbone genes. MGEs from *X. oryzae*, *X. cissicola*, and *X. campestris* were frequently intermingled within the same clusters, indicating extensive sharing of gene content across species boundaries. This pattern suggests horizontal transfer of MGEs or MGE modules within the genus, supporting ongoing or historical exchange among *Xanthomonas* lineages. Although the degree of exchange did vary and some clusters showed enrichment for MGEs from a single species only.

**Fig. 3.**
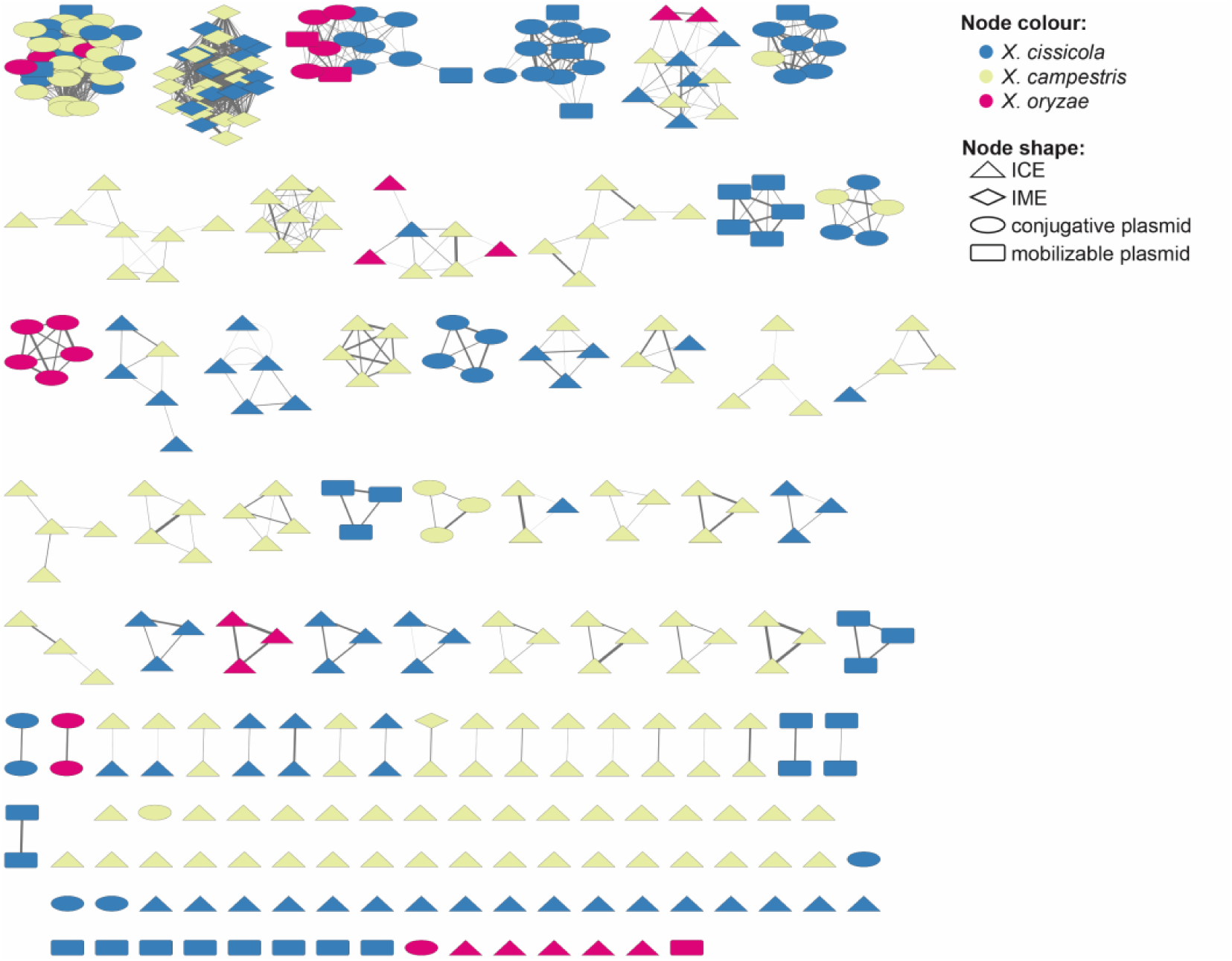
– Gene-content similarity network of MGEs. The network was constructed based on pairwise Jaccard indices calculated from MGE gene content, with edges representing similarities above 0.3. Nodes correspond to individual MGEs and are shaped according to MGE type: IMEs, triangles; ICEs, diamonds; mobilizable plasmids, rectangles; and conjugative plasmids, ellipses. Node colors indicate host species: *X. oryzae* (pink), *X. cissicola* (blue), and *X. campestris* (yellow).

Next, we examined the distribution of the identified MGEs within and between the analysed *Xanthomonas* species. Although no evidence of MGE movement was detected among *X. oryzae* genomes, multiple instances of element sharing were observed among *X. campestris* and *X. cissicola* genomes. One ICE exhibited inter-species movement, being detected in *X. campestris* pv. *barbareae* (GCA_028750075.1) and in two *X. cissicola* pv. *glycines* strains (GCA_001854145.2 and GCA_007567665.1). This ICE encodes a T6SS, two anti-phage defence systems, and two T3SEs (*xopH1* and *avrBs1*) (Fig. 4). In *X. campestris*, only one plasmid, which carries a CALIN element (Cluster of *attC* sites lacking an integron-integrase), was shared between pathovars, being present in one pv. *campestris* genome and in two pv. *incanae* genomes (Fig. S1 and S4). In addition, a *X. cissicola* ICE encoding a T6SS, two anti-phage defence systems, and the T3SE *xopE2* and *xopE3* was identified in one pv. *glycines* strain and in two pv. *fuscans* strains belonging to the fuscans and GL3 lineages. An *X. cissicola* pv. *mangiferaeindicae* genome also harboured an ICE that was also present in both pv. *citri* and pv. *citri* AwA strains. In this same *X. cissicola* pv. *mangiferaeindicae* genome there was also a small plasmid that was also present in the two pv. *citri* genomes (Fig. S2 and S5). This is the only plasmid in *Xanthomonas* observed to carry integron cassettes [55]. No evidence of IME movement was detected.

**Fig. 4.**
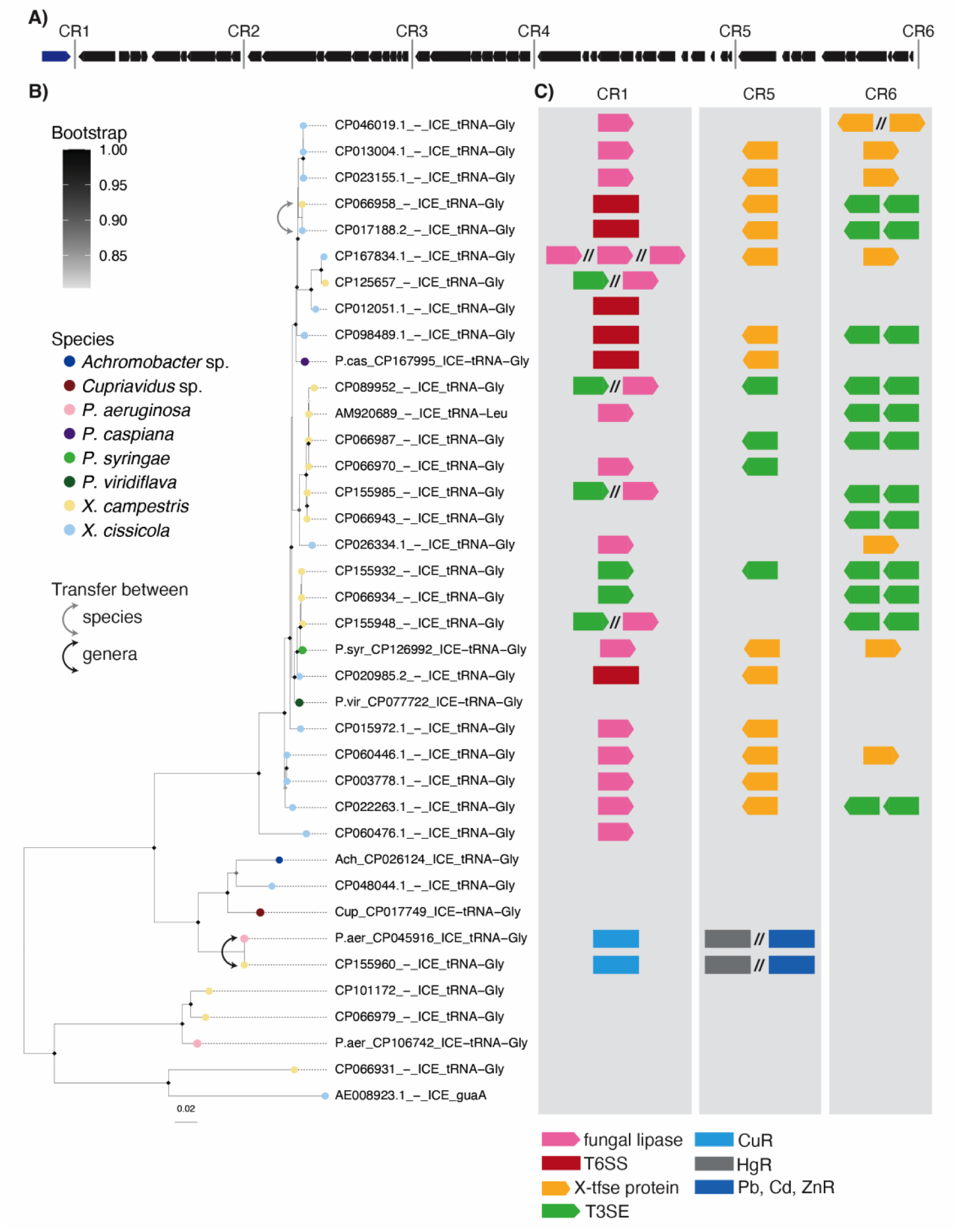
– *Xanthomonas* ICEs. A) Xanthomonas ICEs backbone structure, gene in blue is the site-specific integrase. B) FastTree tree based on the concatenation of alignments of 21 single-copy backbone genes stripped of gaps (21,011 bp). The tree was rooted at midpoint. Diamonds at nodes represent bootstrap support values; only values >80 are shown. Scale bar indicates substitutions per site. C) Representation of relevant cargo genes found in cargo regions (CRs)1, 5 and 6. Not all the cargo genes in the regions are represented. T3SE in CR1 is XopAH, in CR5 is and in CR6 are XopE3 AvrBS1 and XopH1. Genes and operons length are not in scale.

### Conserved backbone architecture and long-range horizontal transfer of *Xanthomonas* ICEs

Overall, ICEs appeared to move across greater phylogenetic distances than plasmids, and the only instance of inter-species MGE movement detected in the *Xanthomonas* genomes involved an ICE. To place this ICE family in a broader evolutionary context, we searched for related elements in other bacterial genera. Using BLASTn, we screened complete genomes available in NCBI, excluding *Xanthomonas* (taxid: 32033), and retained hits showing more than 50% sequence coverage and at least 92% nucleotide identity. Genomes matching these criteria were downloaded, and ICEs were extracted when *att* sites could be identified. This approach identified six ICEs harboured by strains belonging to four *Pseudomonas*, one *Achromobacter*, and one *Cupriavidus* species (Table S4). All of these ICEs were integrated adjacent to a tRNA-Gly locus. Of particular note, *Pseudomonas aeruginosa* strain CF39S, a cystic fibrosis patient isolate, carried an ICE that was nearly identical (differing by only eight SNPs across 107,806 bp) to an ICE found in *X. campestris* pv. *raphani* (GCA_045715785.1). This ICE encodes copper and mercury resistance genes. The presence of nearly identical ICEs in two distinct genera indicates a relatively recent horizontal transfer event between plant-associated *Xanthomonas* and clinical *P. aeruginosa*. The same ICE was also detected in other *P. aeruginosa* strains isolated from other human clinical samples and a *Pseudomonas* sp. strain (CP120377) isolated from soil in Svalbard (Norwegian Arctic archipelago).

While the conjugative plasmids in *Xanthomonas* belong to diverse families, carrying different Mob genes and conjugative systems, ICEs form a single family with a conserved set of backbone genes (Fig. 4A, Table S5). ICE sequences typically consist of conserved backbone genes, which mediate ICE maintenance, regulation, and transfer, and variable accessory or cargo genes [16, 18, 56]. Past work has shown *Xanthomonas* ICEs belong the broad family of the ICE*clc* ICEs, firstly identified in *P. knackmussii* B13 [57], and also frequently present in *P. aeruginosa* [58]. *Xanthomonas* ICEs and ICE*clc* share the integration site (tRNA-Gly) and a syntenic backbone, with a single backbone gene pairwise identity ranging from 63.9% and 66.1% (backbone #4) to 97.9% and 96.2% (backbone #18) when compared with CP155960_-_ICE_tRNA-Gly and CP125657_– _ICE_tRNA-Gly respectively. In this study characterization of the *Xanthomonas* ICE backbone was guided by identification of the set of genes present among all ICEs, except one, using Proteinortho [43] followed by sequence alignment. The relaxed criterion chosen here (present in all ICEs except one) reflects the possibility of gene deletion events. For instance, the site-specific integrase, which is essential for ICE movement [59], appears to have been deleted in AM920689_– _ICE_tRNA-Leu, while CP046019.1_-_ICE_tRNA-Gly lacks from backbone gene #36 to #44. Such deletions may render these ICEs non-mobilizable.

Based on this analysis, the *Xanthomonas* ICE backbone comprises 45 genes (∼37 kb), with predicted functions including ICE maintenance, regulation, and transmission, as well as conserved hypothetical proteins (Table S5). Homologs of these genes in other ICEs encode key components such as conjugative pili (tfc), ICE transfer proteins (tra), and partitioning proteins (par). The backbone genes are in synteny across the ICE, with the exception of CP066931_– _ICE_tRNA-Gly (in *X. campestris*) and AE008923.1_-_ICE_guaA (in *X. cissicola*), which harbour *mobH* and *traG* at the end rather than the beginning of the element.

A phylogeny of *Xanthomonas* ICEs was reconstructed using backbone genes conserved across all ICEs, as well as the seven highly similar ICEs identified from other genera (Fig. 4B). ICEs from *X. cissicola* and *X. campestris* did not form strictly separate clusters, indicating historical exchange between these species. Notably, *Xanthomonas* ICEs did not form a monophyletic group relative to ICEs from other genera. For example, CP048044.1*_*ICE_tRNA-Gly is more closely related to an ICE from *Achromobacter* than to other *Xanthomonas* ICEs, while ICEs from *P. viridiflava, P. syringae, and P. caspiana* branch within the main *Xanthomonas* ICE clade. Together with the nearly identical ICE shared between *P. aeruginosa* and *X. campestris* (CP155960*-*_ICE_tRNA-Gly), these observations indicate that this ICE family is capable of horizontal transfer across long phylogenetic distances.

### MGE accessory genes contribute to plant and environment adaptation

The identified MGEs carry approximately 28.3%, 26.7% and 3.6% of the identified pangenome of *X. campestris*, *X. cissicola* and *X. oryzae*, respectively. Sequence-based homology searches against eggNOG 5.0 [39] revealed that proteins of unknown function are the most abundant category in MGEs (Fig. S6), a trend consistent with prior reports [16, 60]. Analysis of proteins assigned to other categories showed *X. campestris* and *X. cissicola* MGEs were enriched (Fisher’s Exact Test, *p* < 0.01) in proteins involved in replication, recombination, and repair. This category includes transposases and insertion sequences (IS), both known to be common in MGEs [60, 61]. This enrichment suggests MGEs act as hot spots for transposon insertion. Interestingly, *X. oryzae* genomes exhibit a higher proportion of proteins associated with replication/recombination/repair (20.6%) compared to *X. campestris* (6.9%) and *X. cissicola* (7.7%), suggesting that species-specific genomic architectures may influence the potential for generating functional diversity.

MGEs, especially ICEs and IMEs, have recently been revealed to carry a vast repertoire of defence systems (DS) against phage infections [60, 62]. We used DefenseFinder [44] to evaluate the presence and distribution of these genes across MGEs (Fig. 5 and S7, S8, S9). In *X. cissicola*, DS were detected in 44.6% of plasmids 68.7% of ICEs, and 66.7% of IMEs. In *X. campestris*, 37% of plasmids, all but one ICE, and 53.8% of IMEs carried at least one DS. In *X. oryzae*, 33.3% plasmids carried a DS, as did almost all IMEs (9 of 11; 81.8%). In our dataset MGEs harboured up to five different DSs within a single element.

**Fig. 5.**
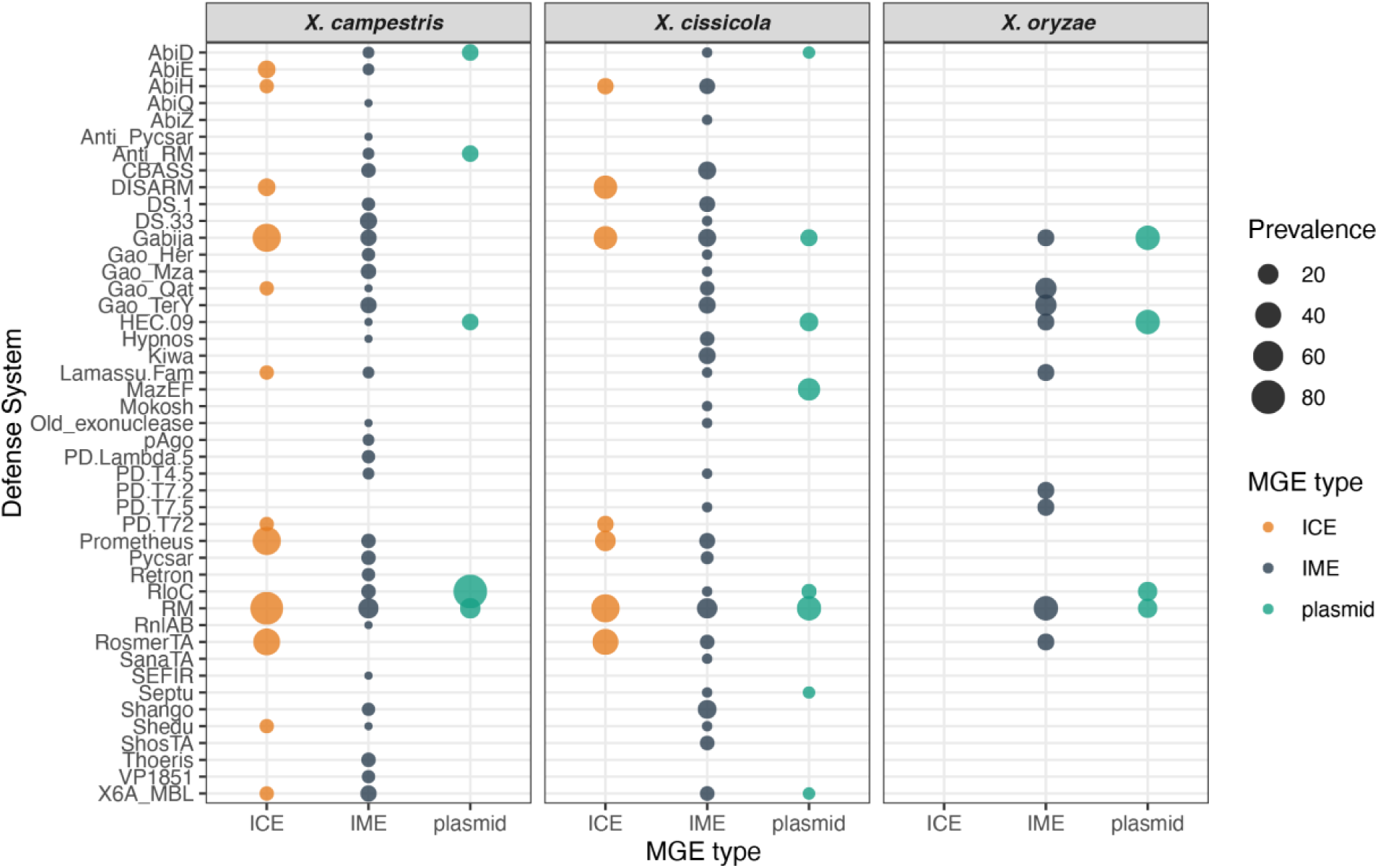
Defence systems in Xanthomonas MGEs. Distribution of defense system genes across X*. campestris*, *X. cissicola* and *X. oryzae* MGEs. The size of the circles is proportional to the percentage of a particular defense system per MGE type per species.

A recent study reported that ICEs generally accumulate more defence systems than IMEs in bacterial genomes [60]. However, in our dataset, IMEs exhibited a more variable repertoire of DS, with ICEs carrying 12 and 9 distinct DS family in *X. campestris* and *X. cissicola*, respectively, whereas IMEs carried 31 and 26 different DS family in the same species. Beyond simple prevalence, the number of defense systems per element was significantly influenced by the type of MGE (Kruskal-Wallis test, *x^2^* = 43.56, *df* = 2, *p* = 3.48 10^−10^), indicating a non-random distribution of these protective suites across the mobilome. Although IMEs were the most prevalent MGE in our dataset, their increased diversity was not merely a sampling artifact. Rarefaction analysis (Fig. S10) demonstrated that IMEs possess a significantly more variable repertoire of DS families compared to both ICEs and plasmids. Even when normalized to an equivalent sampling depth, IMEs exhibited a steeper accumulation rate and higher richness of unique DS families (e.g., reaching over 40 distinct families) compared to ICEs and plasmids, which showed early saturation. Across all three species, the most prevalent DSs were restriction–modification (RM) systems, followed by Gabija systems (Fig. 5). This is in keeping with other analyses of DSs in prokaryotic genomes, with RM systems reported as most common, present in approximately 80% of genomes, with Gabija systems which degrade phage and bacterial genomic DNA upon phage infection [63], also reported as widespread, occurring in roughly 20% of genomes [44].

Type three secretion system effectors (T3SEs) are known to be key determinants of host interaction and virulence in phytopathogens, to identify T3SEs, we used a database of previously characterized T3SEs retrieved from the EuroXanth platform [42, 64]. In *X. campestris*, the majority of ICEs (all except four) carried at least one T3SE, all of which were either XopH1, XopAH, XopE3, or AvrBs1 (Fig. 4, S11 and S12). In plasmids of this same species there was lower proportion showing carriage, with T3SEs observed in 59% of plasmids but the effector repertoire was more diverse (XopJ6, AvrBs3 (TALE), XopAH, XopJ5, XopAF2, XopE2, XopAA, XopAE, and XopAO) (Fig. S4 and S12). Among the 117 non-redundant IMEs, only ten carried a T3SE, seven of which encoded XopG1 (Fig. S12). In *X. cissicola*, T3SEs were comparatively rare on ICEs but were frequently encoded on plasmids (Fig. 4, S5 and S12). In these genomes 63% of plasmids carry at least one effector and some plasmids encode up to six T3SEs with considerable diversity of type, including AvrBs3, XopE3, XopG1, XopAH, XopAF2, XopC1, XopJ1, XopJ4, XopJ5, and XopT (Figs. S5 and S12).

Carriage of antimicrobial resistance (AMR) determinants by MGEs is also of high interest, with these elements often responsible for AMR gene spread among prokaryotes. While in crop production, clinically-relevant antibiotics are not generally used, other antimicrobial agents, most commonly copper [23], are often applied, both in conventional and organic farming [65]. A number of past studies have shown MGEs in *Xanthomonas* and other plant pathogens carry copper, arsenic resistance genes [12, 25, 66, 67] with antibiotic resistance genes also recovered in some instances [68]. Carriage of copper and other heavy metal resistance genes was observed in relatively few MGEs from the *Xanthomonas* species analysed here. In *X. oryzae*, a conjugative plasmid carried the *czc* operon, which confers resistance to cadmium, zinc, and cobalt [69], while a mobilizable plasmid encoded arsenic resistance genes. In *X. campestris*, one plasmid encoded copper resistance and one ICE, sharing high homology to an ICE found in *P. aeruginosa* strain CF39S harboured copper, lead, zinc and cadmium, and mercury resistance genes (Fig. 4). In *X. cissicola*, three different plasmids contained arsenic and copper resistance genes as well as the *czc* operon, with an additional plasmid containing copper resistance genes and the *czc* operon (Fig. S5).

Pangenome association analysis was performed to assess whether particular MGE cargo genes were associated with specific pathovars. In *X. campestris*, the effector gene, *xopAH*, was significantly associated with pv. *campestris*, and was introduced into this pathovar via ICEs. Notably, *xopAH*, and the other effector genes encoded by *campestris* ICEs (*avrBs1*, *xopH1, xopE3*) were previously reported as horizontally transferred, however, their mobility was attributed to nearby transposases, and the ICEs carrying these genes were not identified [54]. Also in *X. campestris,* the pv. *incanae* representatives were significantly associated with a gene encoding an X-Tfes protein containing a XVIPCD domain, a conserved ∼120-amino acid C-terminal region characteristic of X-Tfes toxins. These toxins belong to a family of bacterial effectors in *Xanthomonas* species known to mediate interbacterial antagonism via the T4SS [70]. This gene was also frequently detected in *X. cissicola* ICEs, being present in 11 non-redundant ICEs, with four ICEs carrying up to four copies of the gene (Fig 4C).

In *X. cissicola*, a TALE effector encoding gene (*avrBs3*) carried on a mobilizable plasmid was significantly associated with pv. *fuscans*, a pathovar for which there are three distinct lineages (GL2, GL3, and *fuscans*) that are not monophyletic. Notably, pv. *aurantifolii*, which causes citrus canker, phylogenetically clusters within the *fuscans* complex but lacks this plasmid (Fig. S2). This pattern suggests that acquisition of the plasmid carrying *avrBs3* may have contributed to the adaptation of pv. *fuscans* strains to bean hosts, rather than reflecting shared ancestry. In pv. *glycines*, multiple genes encoded by an IME integrated at a tRNA-Arg locus were significantly associated with the pathovar, including two of the four defence systems carried by this element (Gao_Qat and CBASS).

To identify genes shared across MGEs that may contribute to plant or environmental adaptation we compared the predicted proteins encoded by these elements using Proteinortho. Eight of 15 campestris ICEs and 12 of 16 cissicola ICEs were found to carry a gene containing a fungal lipase-type domain (Fig. 4C). This gene may encode an enzyme involved in the degradation of plant cell wall components, suggesting a potential role in host interaction. This also revealed that a type VI secretion system (T6SS), likely to mediate interactions with other microbes, was carried by four

*X. campestris* and one *X. cissicola* ICEs (Fig. 4C). EcoFoldDB-annotate [71], which identifies structural homologs of proteins key for biogeochemical and plant-microbe interaction traits, identified a pectate liase in four *X. cissicola* plasmids present in almost all pv. *glycinae* genomes, and in *X. campestris*. Pectate lyase is an enzyme that degrades plant cell walls by cleaving pectin, and in *X. cissicola*, a pectate lyase homologue was shown to be associated with the water-soaked margins of canker lesions [72]. To identify secondary metabolites, we utilized antiSMASH 8.0 [73] which revealed a *X. cissicola* IME carrying a gene cluster involved in synthesizing kolossin, a secondary metabolite hypothesized to be involved in interspecies communication [74].

The ICE AE008923.1_-_ICE_guaA was detected in all pv. *citri* A strains of *X. cissicola*, although in a fragmented form and therefore not included in the ICE repertoire analysed here. This element was also present in all pv. *citri* AwA strains, being intact in all but one genome, and was detected in either functional or degraded forms across all pv. *citri* strains examined. These observations suggest that this ICE was acquired prior to pv. *citri* diversification and may have contributed to the expansion of this pathovar. However, this element does not appear to encode genes directly implicated in plant adaptation; instead, it carries two distinct defence systems, Lamassu and Septu [75].

## DISCUSSION

Across bacteria, MGEs are a major contributor to rapid adaptation, yet closely related lineages can differ substantially in the elements and associated gene cargo they contain. Here, we present a comprehensive comparative analysis of conjugative and mobilizable plasmids, ICEs, and IMEs across three major species of *Xanthomonas*. We find pronounced heterogeneity in MGE content:

*X. cissicola* and *X. campestris* harbour large and diverse MGE repertoires, whereas *X. oryzae* contains remarkably few mobile elements. Host defence provides a partial explanation for the differences in MGE carriage across *Xanthomonas* species. In *X. campestris*, CRISPR-Cas presence is clearly associated with reduced MGE load, consistent with previous report linking loss of CRISPR-Cas immunity to enhanced genome plasticity and diversification of the pv. *campestris* [54]. In contrast, CRISPR-Cas immunity appears comparatively ineffective against MGEs in *X. cissicola,* with the low number of spacers per array supporting the idea that MGEs in *X. cissicola* may be routinely bypassing CRISPR-Cas interference, rather than being excluded. In *X. oryzae*, the presence of CRISPR-Cas does not explain the scarcity of MGEs, suggesting that other defense systems such as restriction-modification systems contribute to defense from MGE invasion [76]. Notably, *X. oryzae* genomes are enriched in transposases and IS relative to *X. cissicola* and *X. campestris*, and have been reported to carry among the highest number of insertion sequences per genome across plant pathogens, with evidence of transposase-driven gene disruption, recombination, and dissemination of fitness-associated genes [77]. A recent study showed that recombination has a huge impact in the evolution of the Asian-like strains of *X. oryzae* pv*. oryzae* [78]. This suggests that these species within the same genus have different evolutionary strategies, *X. cissicola* and *X. campestris* innovate via MGE accumulation, *X. oryzae* via homologous recombination and rearrangements.

ICEs in *Xanthomonas* belong the broad family of the ICE*clc* ICEs. Phylogenetic and network analyses reveal that these ICEs are not confined to a single species or even genus, the discovery of nearly identical ICEs in *Xanthomonas* and two *Psudomonas* species, provides evidence for recent horizontal transfer across large phylogenetic distances. The fact that almost identical ICEs are found across plant-associated *Xanthomonas*, soil-isolated *Pseudomonas* sp., and clinical isolates of *P. aeruginosa* strongly suggests a multistep horizontal transfer linking agricultural, environmental, and clinical ecosystems. Phylogenetic and network analyses reveal that these *Xanthomonas* ICEs belong to the broad ICEclc family and are not confined to a single species or genus. In fact, the discovery of nearly identical elements across such large phylogenetic distances provides evidence of recent horizontal gene transfer. As these elements move through different environments, their cargo adapts: while ICEclc-type elements in clinical P. aeruginosa isolates are enriched for heavy metal resistance, potential efflux systems, and multidrug resistance proteins [58], in *Xanthomonas*, only one ICE carried antimicrobial resistance. Although copper resistance is environmental relevant trait for a plant pathogen, in *Xanthomonas*, copper resistance is more common on plasmids,with ICEs carrying other environmental-relevant traits. This genus-spanning variation in cargo genes mirrors the SXT-R391 ICE family, where clinical variants typically carry antibiotic resistance [79], while variants from free-living marine Alteromonas encode metal resistance and restriction-modification systems [80].

Overall, our analyses demonstrate that in many *Xanthomonas* MGEs are key vehicles for the dissemination of genes involved in host interaction, virulence, and environmental adaptation. Most notably, we detected a considerable number of T3SEs embedded within MGEs, with plasmids carrying a highly variable repertoire of these key determinants of host interaction and virulence. Additionally, the association of specific effectors with particular pathovars, such as *xopAH* in pv. *campestris* and TALE (*avrBs3*) in pv. *fuscans*, along with the frequent carriage of genes encoding putative plant cell wall-degrading enzymes, such as pectate lyases and fungal lipase-like proteins, supports a model in which MGE acquisition contributes to host specialization. T3SEs were frequently found to co-occur with defense system genes across MGEs. In contrast, T3SEs were never detected on the same MGE as heavy metal resistance genes, pointing to a functional specialization of individual elements. This pattern suggests that certain adaptive functions may be selectively compatible when co-localized on the same element, whereas others may be antagonistic or impose fitness trade-offs that disfavor their coexistence. These observations are partially consistent with a recent large-scale analysis of ICEs and IMEs across prokaryotic genomes, which reported negative correlations between antimicrobial resistance, virulence genes, but also defense systems, within individual mobile elements [60]. MGEs in *Xanthomonas* may tend to specialize in distinct functional roles, however, host genomes can accommodate this specialization by accumulating multiple MGEs with complementary adaptive functions. Interbacterial competition and defence in *Xanthomonas* may also be linked to genes present within MGEs. The frequent occurrence of X-Tfes toxins, defence systems including restriction-modification, Gabija and CBASS, and secretion system components on ICEs and IMEs suggests that these elements mediate not only adaptation to plant hosts but also survival within complex microbial communities.

Similarly to other genera [5, 60, 81], IMEs vastly outnumbered ICEs and plasmids. Contrary to recent reports suggesting that ICEs are the primary carriers of defence systems [60], our data reveal that IMEs often encode a more diverse set of defence families. Although ICEs have been highlighted as major carriers of defence systems [60, 62], this pattern does not appear to be universal across all ICE families. For example, the PsICE family identified in plant-associated *Pseudomonas* species typically lacks defence systems, and only rarely encodes T3SEs [18], indicating that the functional repertoire of ICEs can vary substantially. The high diversity of integration sites, variable gene content, and broad range of defence systems carried by IMEs suggest that these elements may play an important role in environmental adaptation, beyond their traditionally underappreciated contribution to HGT. In this context, although phage-mediated biocontrol is increasingly explored as a strategy for plant disease management [82–84], the widespread presence of anti-phage defence systems within MGEs, particularly IMEs, highlights the potential for rapid dissemination of phage resistance traits within bacterial populations, which may ultimately influence the long-term efficacy and evolutionary stability of such control approaches.

## CONCLUSION

Overall, our study highlights MGEs as key drivers of adaptive evolution in *Xanthomonas* and indicates that the capacity for mobilome expansion is strongly influenced by the diversity and efficacy of host CRISPR-Cas defences. Differences in MGE composition and mobility have contributed to divergent evolutionary trajectories across species, influencing host range, virulence repertoires, and ecological success. By revealing extensive intra– and interspecies MGE exchange and uncovering a globally distributed ICE family, this work underscores the importance of considering MGEs as dynamic and interconnected components of bacterial evolution in agricultural and natural ecosystems.

