## Supplementary Figures for "Diverse conjugative and mobilisable elements underpin key adaptive traits in *Xanthomonas*"

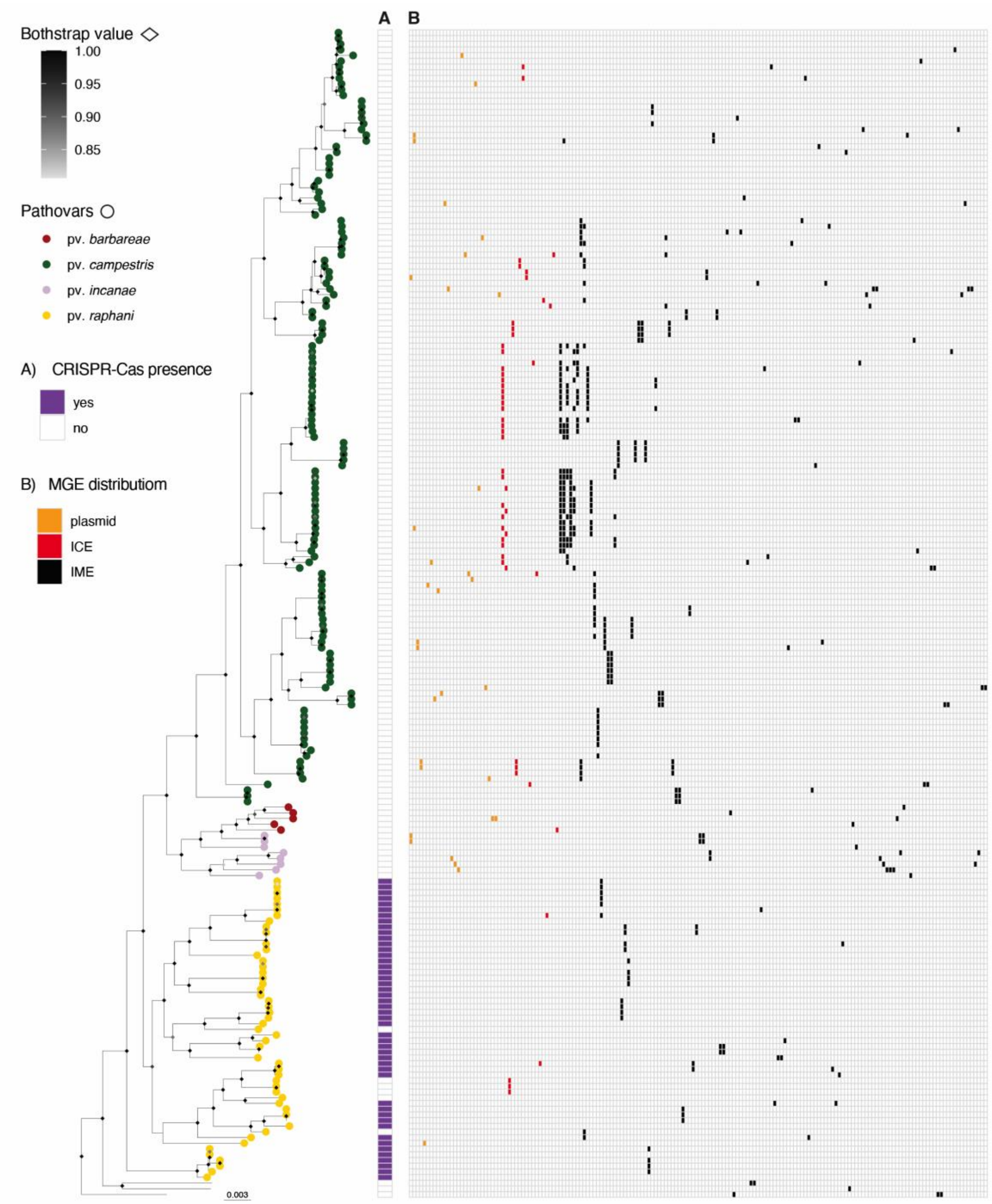


**Fig. S1 - MGEs distribution in *X. campestris.*** The phylogeny of *X. campestris* was built using Realphy, using GCA_000007145.1 as a reference genome. The tree was rooted with GCA_000972745 (*X. arboricola*) then the tip was removed from the tree. Diamond at nodes represent bootstrap support values; only values >80 are shown. Tip labels show pathovars, which were assigned from the literature search (Table S1). The scale bar indicates substitutions per site. A) Indicates the presence of a CRISPR-Cas system. Panel B) represents the distribution of nonredundant MGEs carried by the corresponding isolate. Per each MGE category, MGEs are listed in decreasing order based on the number of strains within the species that carry them.

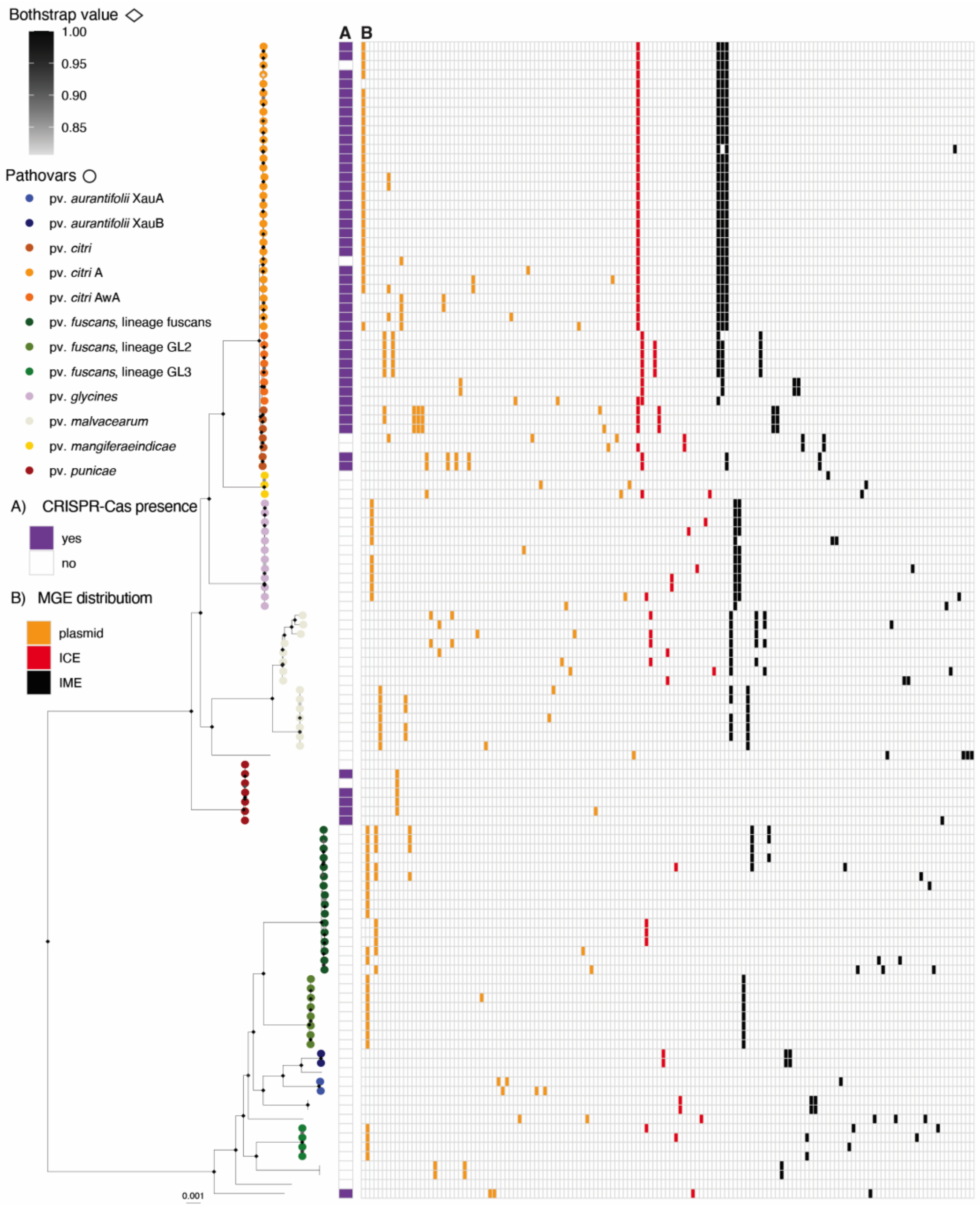


**Fig. S2 - MGEs distribution in *X. cissicola.*** The phylogeny of *X. cissicola* was built using Realphy, using GCA_000007165.1 as a reference genome. The tree was rooted with GCA_000009165.1 (*X. euvesicatoria*) then the tip was removed from the tree. Tip labels show pathovars, which were assigned via literature search (Table S1). Diamond at nodes represent bootstrap support values; only values >80 are shown. Tip labels show pathovars, which were assigned from the literature search. The scale bar indicates substitutions per site. A) Indicates the presence of a CRISPR-Cas system. Panel B) represents the distribution of nonredundant MGEs carried by the corresponding isolate. Per each MGE category, MGEs are listed in decreasing order based on the number of strains within the species that carry them.

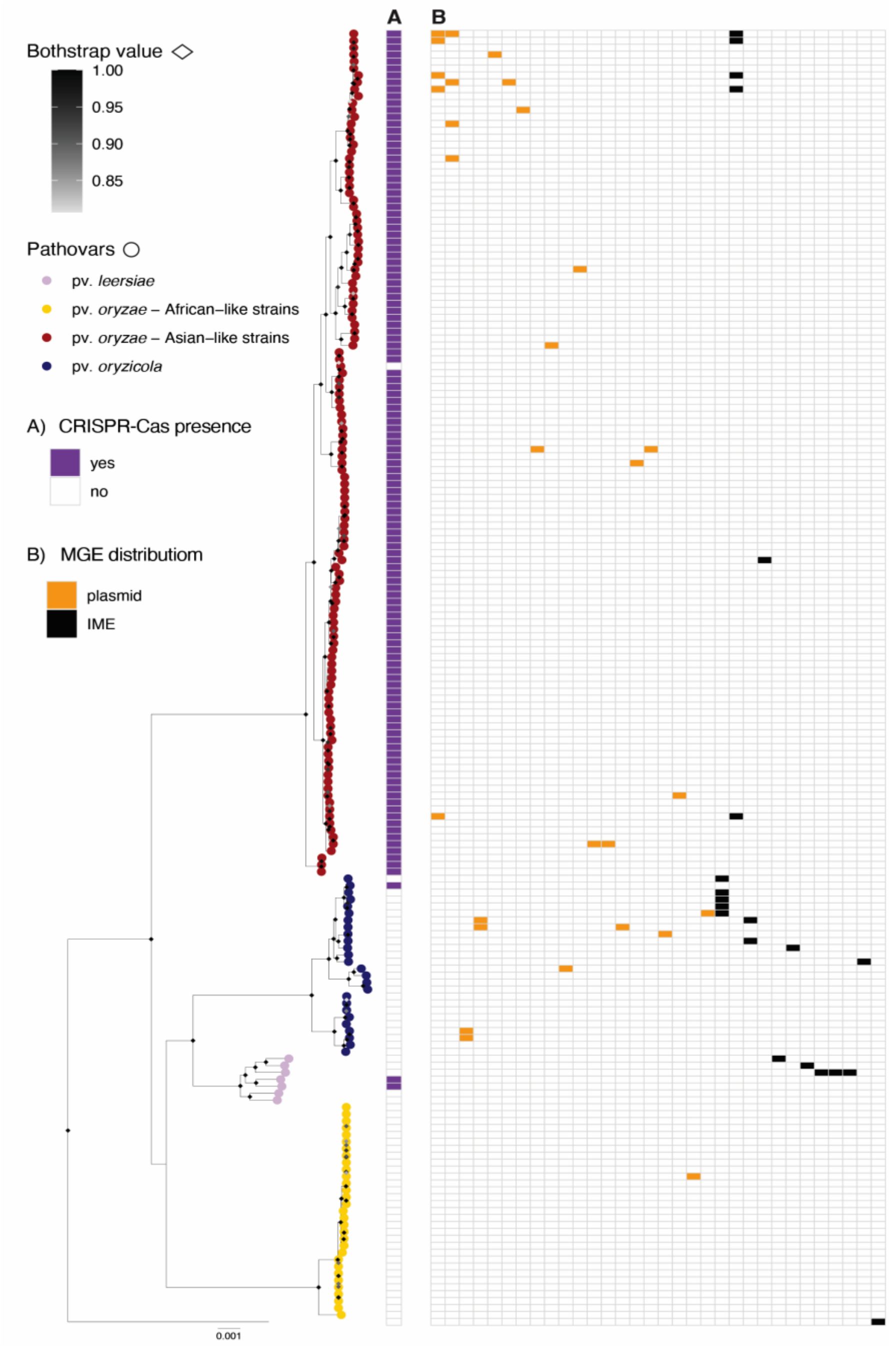


**Fig. S3 - MGEs distribution in *X. oryzae.*** The phylogeny of *X. oryzae* was built using Realphy, using GCA_000007145.1 as a reference genome. The tree was rooted with GCA_002759095.2 (*X. phaseoli*) then the tip removed from the tree. Tip labels show pathovars, which were assigned via literature search (Table S1). Diamond at nodes represent bootstrap support values; only values >80 are shown. Tip labels show pathovars, which were assigned from the literature search. The scale bar indicates substitutions per site. A) Indicates the presence of a CRISPR-Cas system. Panel B) represents the distribution of nonredundant MGEs carried by the corresponding isolate. Per each MGE category, MGEs are listed in decreasing order based on the number of strains within the species that carry them.

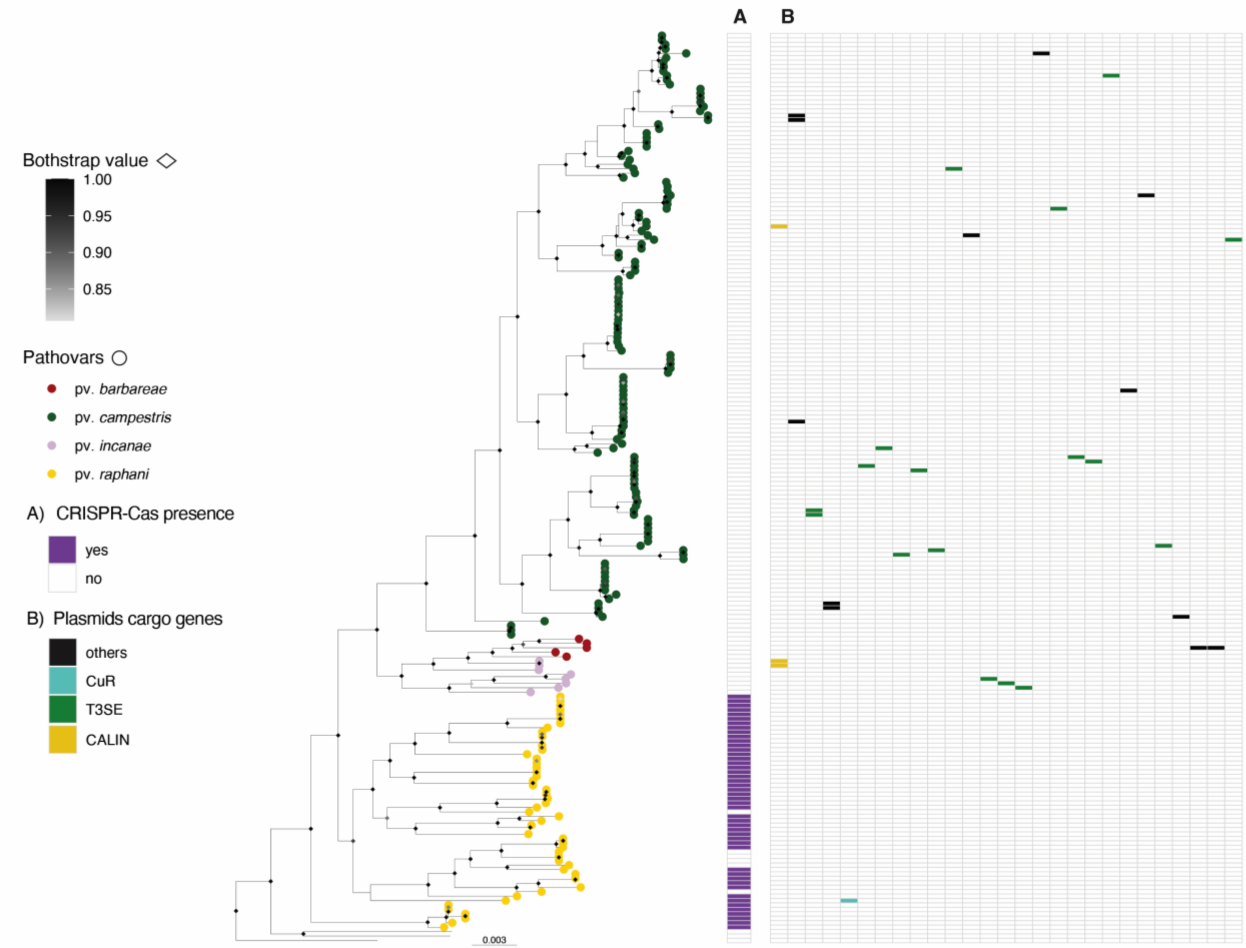


**Fig. S4 - Plasmids distribution in *X. campestris*.** The phylogeny of *X. campestris* was built using Realphy, using GCA_000007145.1 as a reference genome. The tree was rooted with GCA_000972745 (*X. arboricola*) then the tip was removed from the tree. Diamond at nodes represent bootstrap support values; only values >80 are shown. Tip labels show pathovars, which were assigned from the literature search (Table S1). The scale bar indicates substitutions per site. A) Indicates the presence of a CRISPR-Cas system. Panel B) represents the distribution of non-redundant plasmids carried by the corresponding isolate. Non-redundant plasmids are listed in decreasing order based on the number of strains within the species that carry them, colors within B) represent cargo genes category of interest.

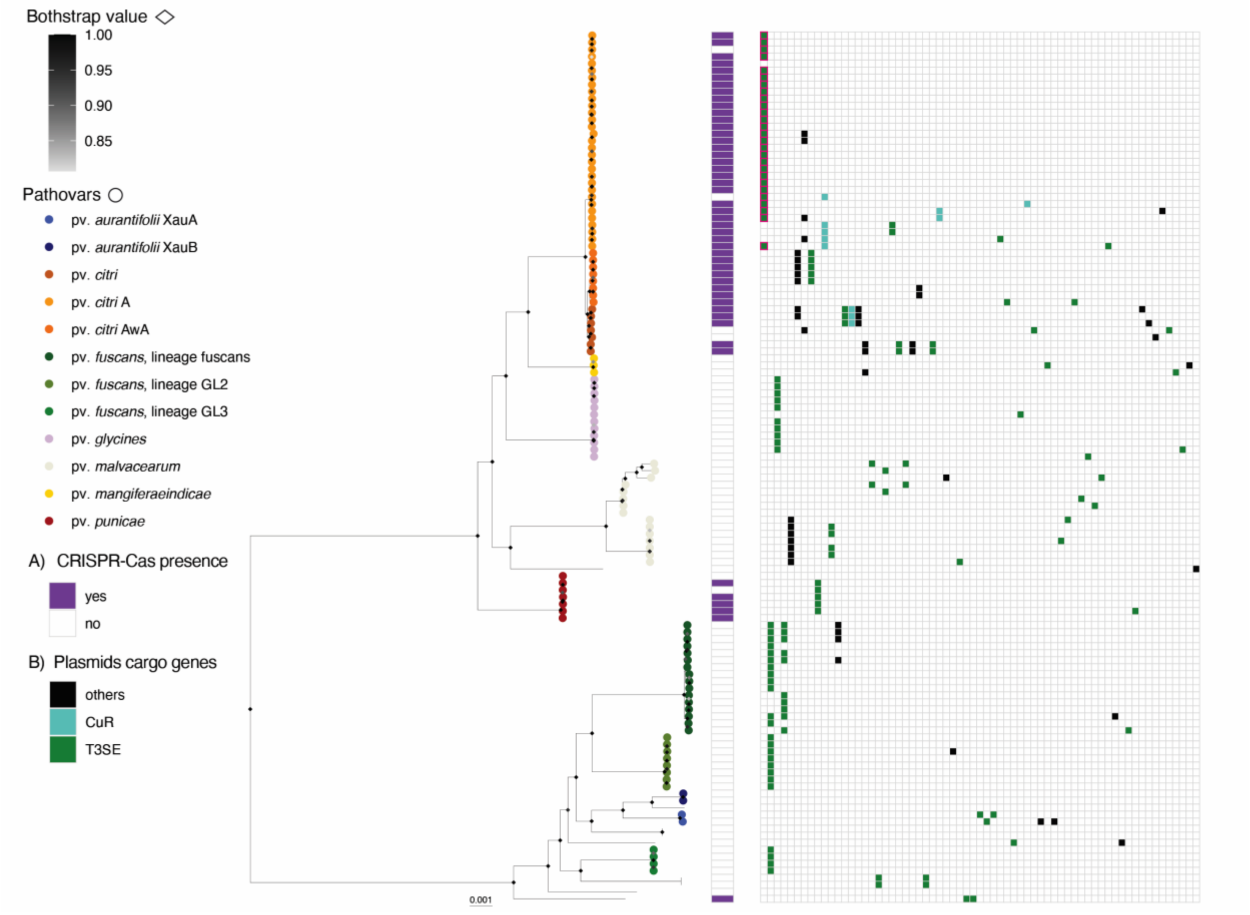


**Fig. S5 - Plasmids distribution in *X. cissicola.*** The phylogeny of *X. cissicola* was built using Realphy, using GCA_000007165.1 as a reference genome. The tree was rooted with GCA_000009165.1 (*X. euvesicatoria*) then the tip was removed from the tree. Tip labels show pathovars, which were assigned via literature search (Table S1). Diamond at nodes represent bootstrap support values; only values >80 are shown. Tip labels show pathovars, which were assigned from the literature search. The scale bar indicates substitutions per site. A) Indicates the presence of a CRISPR-Cas system. Panel B) represents the distribution of non-redundant plasmids carried by the corresponding isolate. Non-redundant plasmids are listed in decreasing order based on the number of strains within the species that carry them, colors within B) represent cargo genes category of interest.

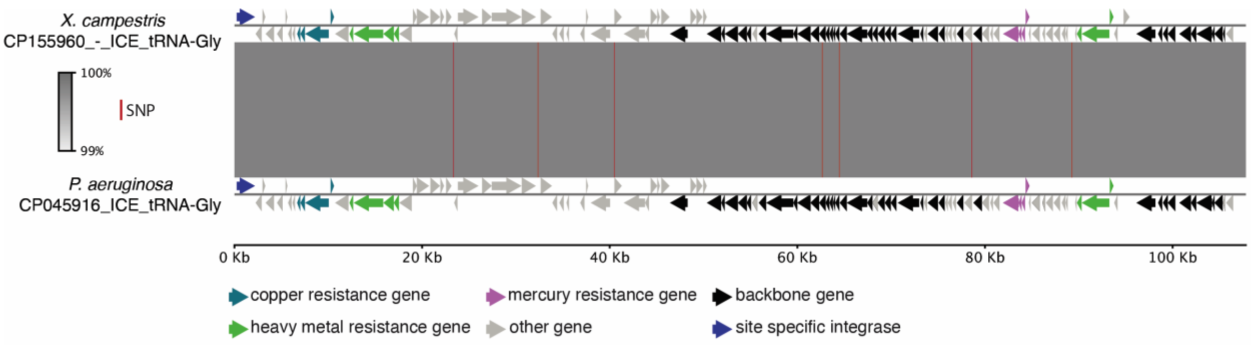


**Fig. S6 – Inter genera ICE movement.** pyGenomeViz (https://github.com/moshi4/pyGenomeViz) alignment between CP155960_-_ICE_tRNA-Gly found in *X. campestris* pv. *raphani* (GCA_045715785.1) and CP045916_ICE_tRNA-Gly isolated from *Pseudomonas aeruginosa* strain CF39S.

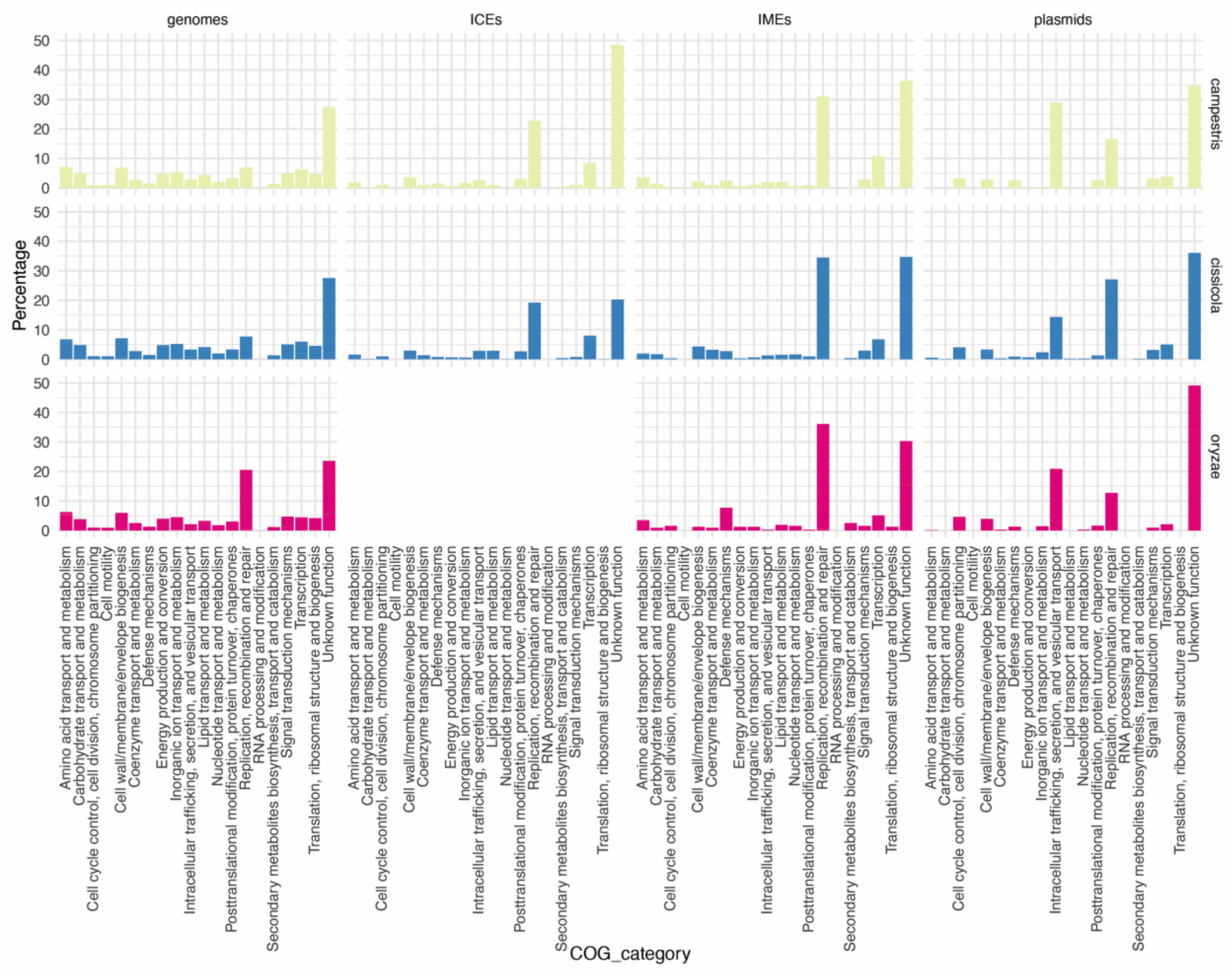


**Fig. S7 - Functional composition of MGEs in *Xanthomonas*.** COG (Clusters of Orthologous Groups) category analysis was performed using eggNOG 5.0. Bars represent the proportion of genes assigned to each COG category for plasmids, ICEs, IMEs, and the corresponding host genomes.

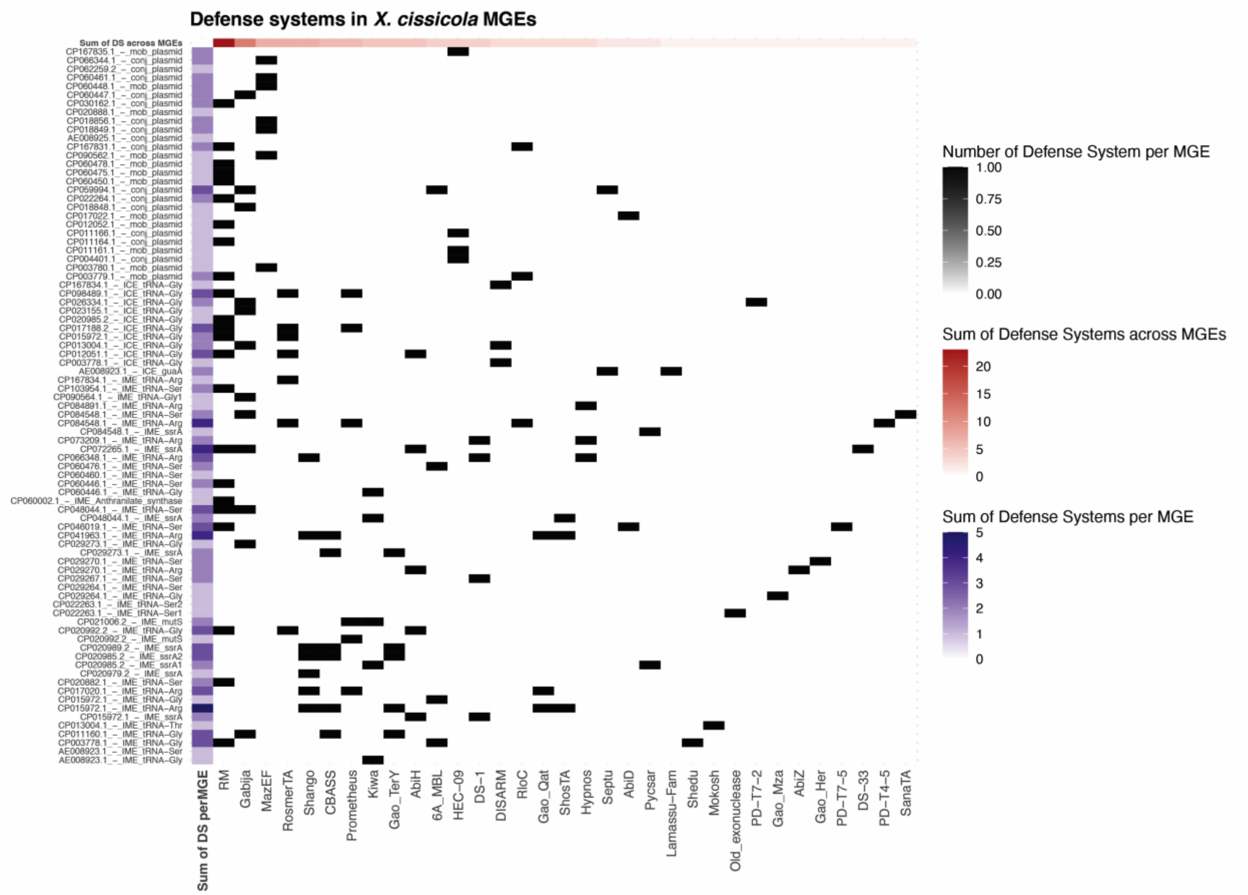


**Fig. S8. Distribution of defence systems across MGEs in *X. cissicola*.** Heatmap showing the abundance of defence systems encoded by different MGEs. Rows correspond to individual MGEs, ordered by MGE type, while columns represent defence system categories, ordered by their cumulative occurrence across MGEs.

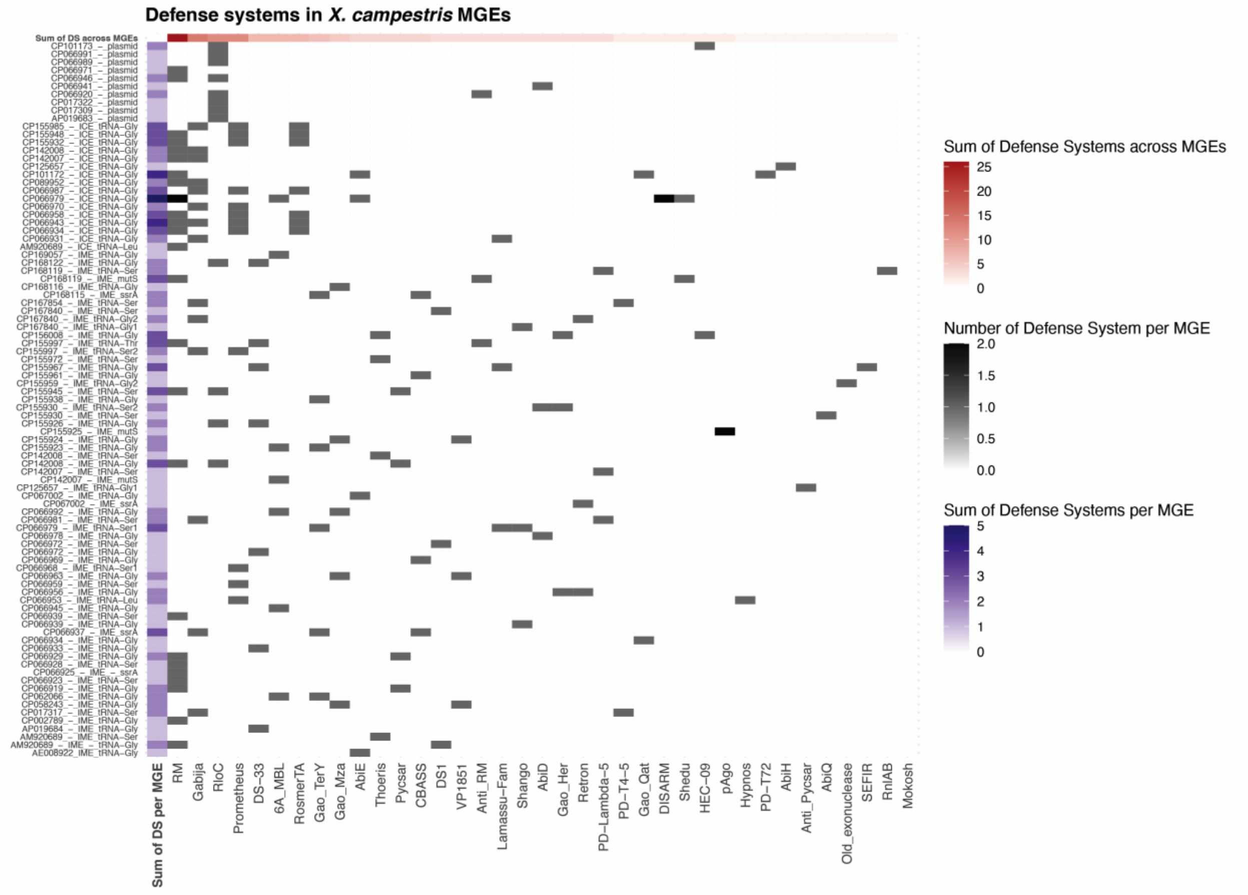


**Fig. S9 - Distribution of defence systems across MGEs in** *X. campestris***.** Heatmap showing the abundance of defence systems encoded by different MGEs. Rows correspond to individual MGEs, ordered by MGE type, while columns represent defence system categories, ordered by their cumulative occurrence across MGEs.

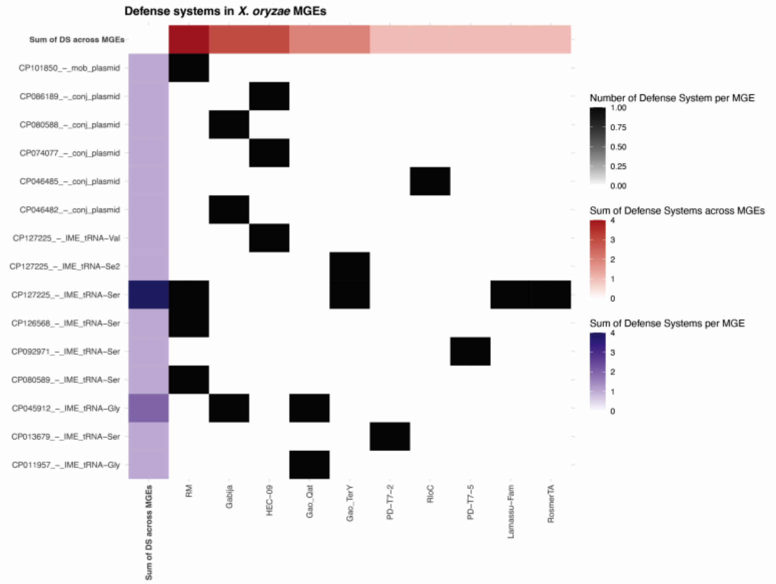

**Fig. S10 - Distribution of defence systems across MGEs in *X. oryzae*.** Heatmap showing the abundance of defence systems encoded by different MGEs. Rows correspond to individual MGEs, ordered by MGE type, while columns represent defence system categories, ordered by their cumulative occurrence across MGEs.

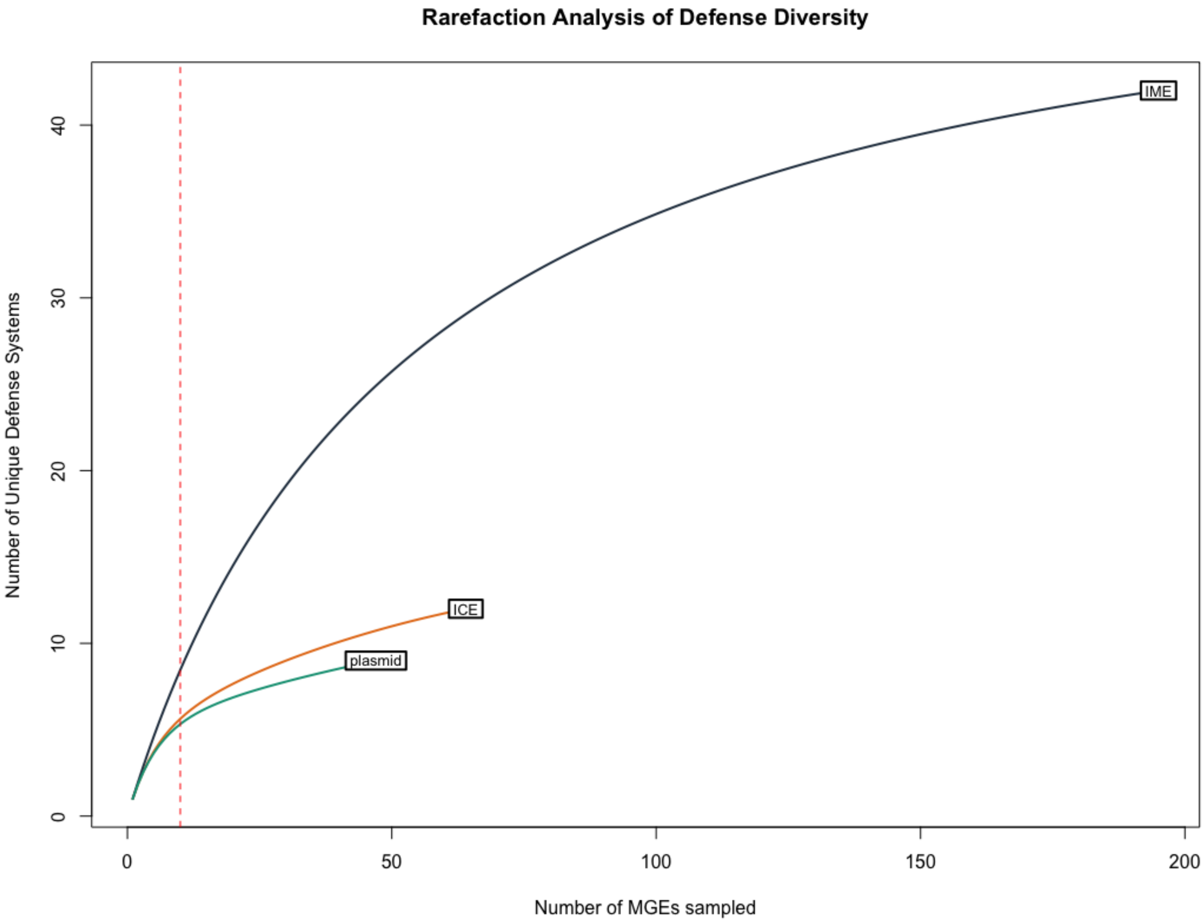


**Fig. S11 - Rarefaction curves of defense system richness.** Comparison of the cumulative number of unique defense system families identified by DefenseFinder across IMEs, ICEs, and plasmids. The steeper slope of the IME curve (dark slate) indicates higher intrinsic diversity independent of sampling effort. Dashed red line represents the rarefaction point for normalized diversity comparisons. Data includes elements from *X. campestris*, *X. cissicola*, and *X. oryzae*.

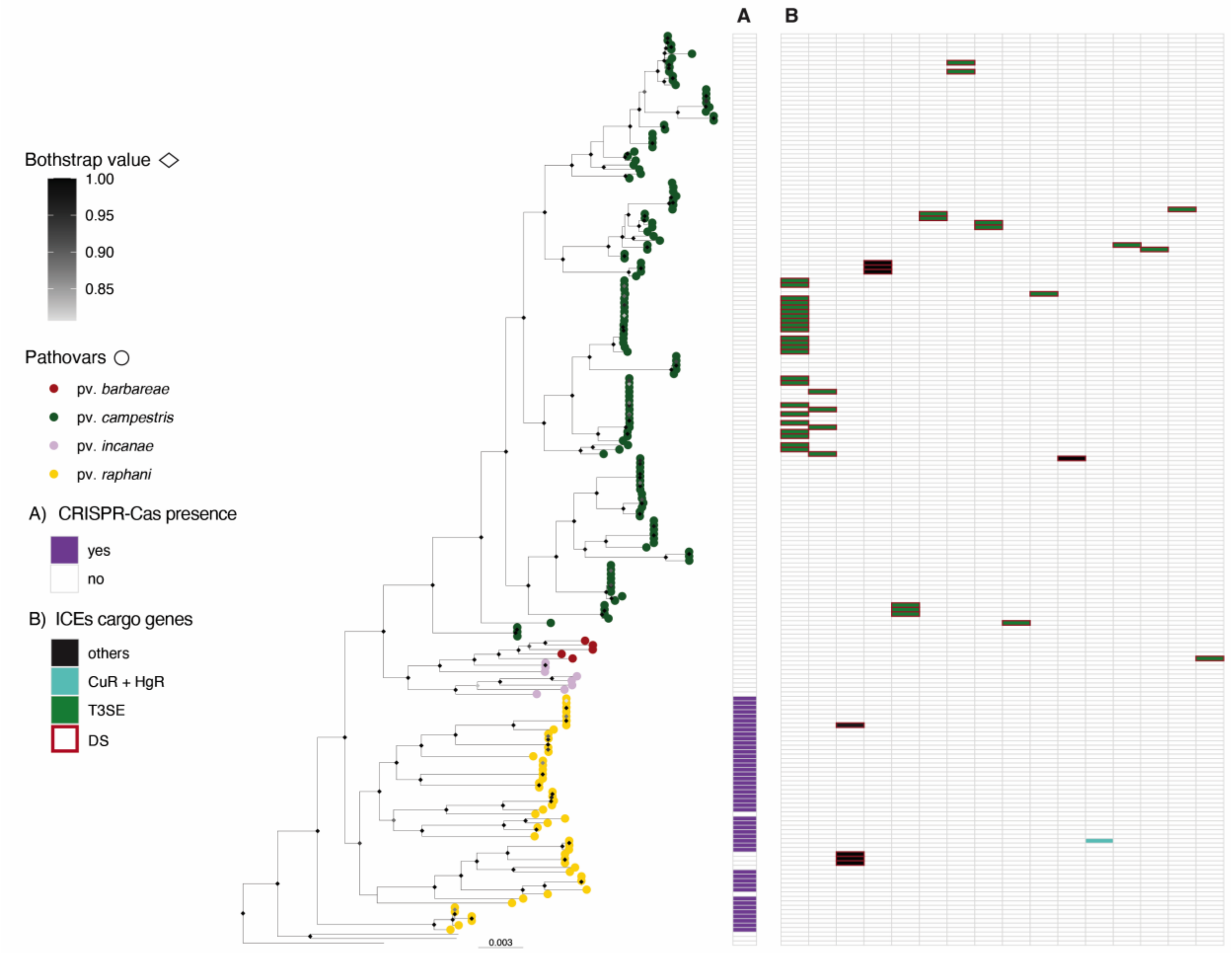


**Fig. S12 - ICE distribution in *X. campestris*.** The phylogeny of *X. campestris* was built using Realphy, using GCA_000007145.1 as a reference genome. The tree was rooted with GCA_000972745 (*X. arboricola*) then the tip was removed from the tree. Diamond at nodes represent bootstrap support values; only values >80 are shown. Tip labels show pathovars, which were assigned from the literature search (Table S1). The scale bar indicates substitutions per site. A) Indicates the presence of a CRISPR-Cas system. Panel B) represents the distribution of non-redundant ICEs carried by the corresponding isolate. Non-redundant ICEs are listed in decreasing order based on the number of strains within the species that carry them, colors within B) represent main cargo genes category.

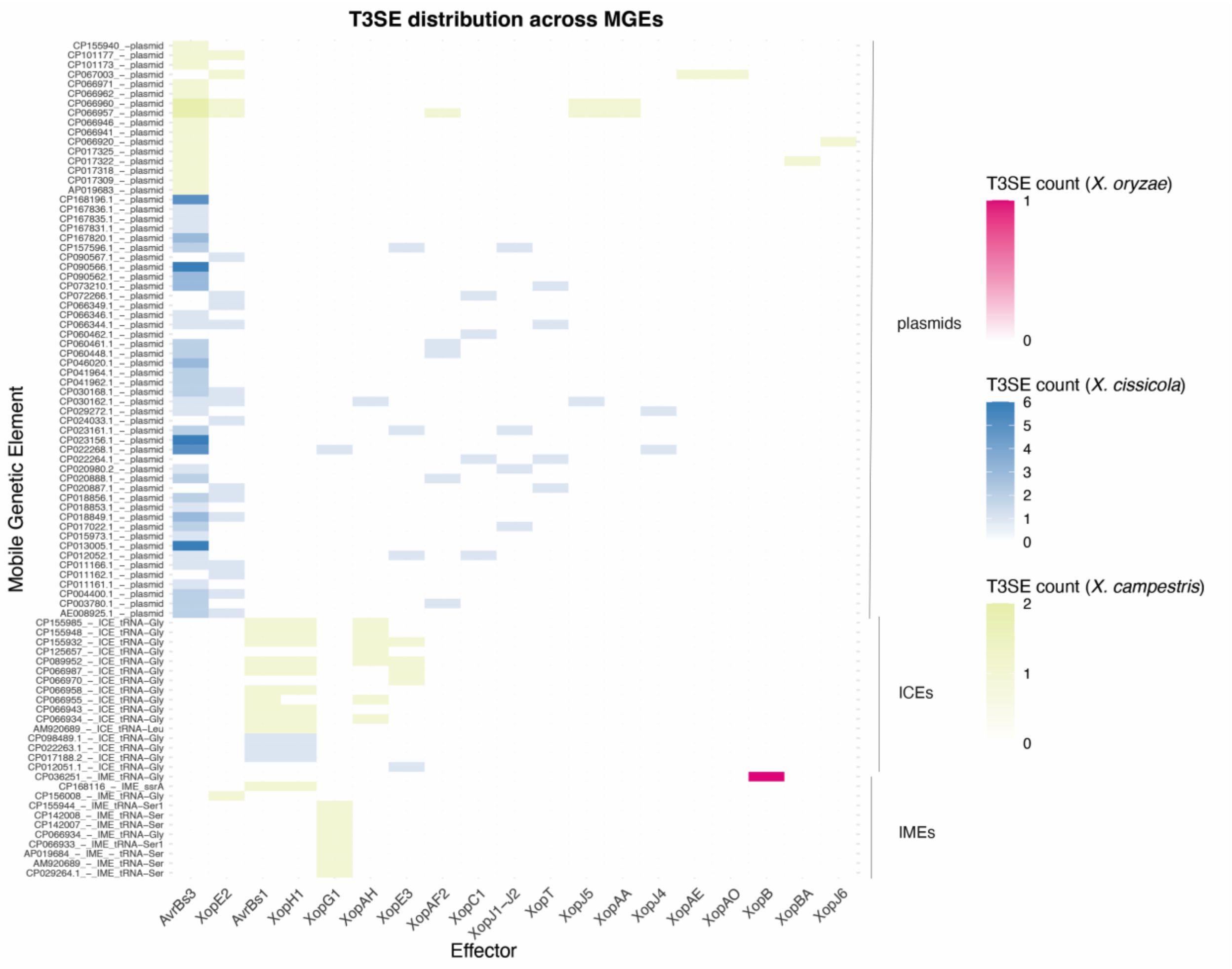


**Fig. S13 - Distribution of T3SE across MGEs.** Heatmap showing the abundance of T3SE encoded by different MGEs. Rows correspond to individual MGEs, ordered by MGE type and by species, while columns represent the different T3SEs, ordered by their cumulative occurrence across MGEs.

NOT IN THE TEXT

Phylogenetic network analysis conducted with SplitsTree of *Xanthomonas* ICE backbone sequences showed moderate phylogenetic conflict (mean delta score = 0.2415; Q-residual = 0.001253), consistent with a largely tree-like signal punctuated by reticulation (Fig. S11). Individual ICE delta scores ranged from 0.14 to 0.37, with several elements exhibiting elevated values (>0.30). Consistent with these patterns, the PHI test detected statistically significant evidence of recombination (p = 0.0), supporting a role of inter-ICE recombination in shaping ICE diversity. In contrast to ICE-associated genes, the core genome exhibited very low individual delta scores (mostly <0.10). This pattern suggests that the increased phylogenetic conflict is specific to ICE-associated regions and arises from their mobile and recombinogenic nature, rather than from the background genomic context.

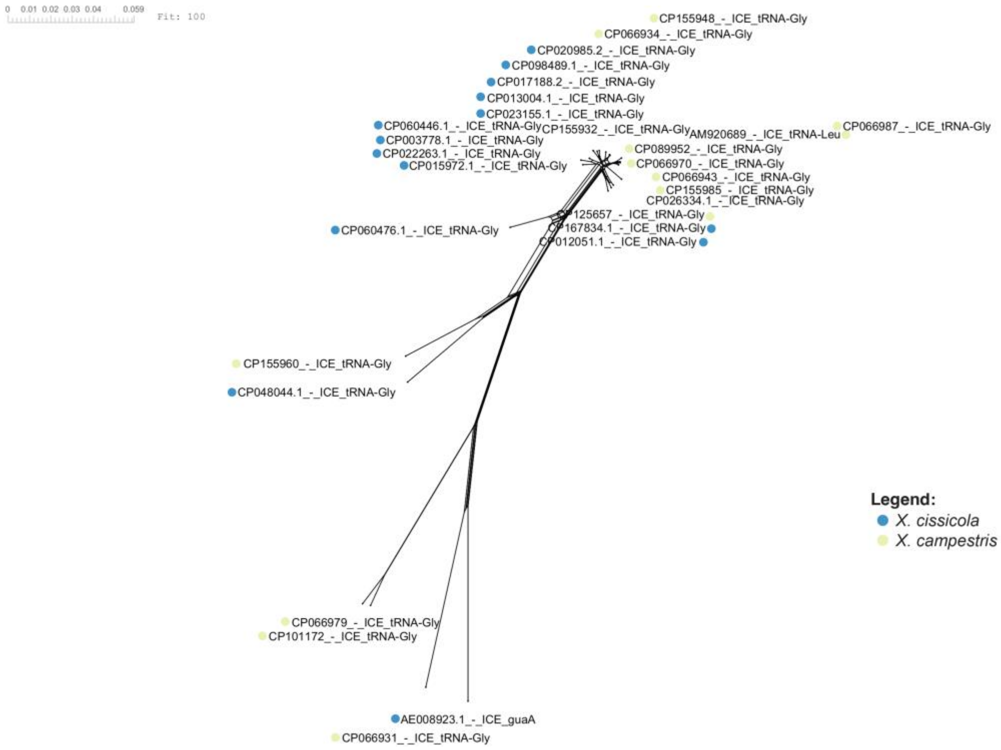


**Fig. S14** – **ICE-core SplitsTree.** Neighbor-Net generated in SplitsTree using a concatenated alignment of backbone genes conserved in all nonredundant ICEs. The scale bar indicates substitutions per site
